# Evolutionary regimes determine the accuracy of epistasis inference from temporal genetic data

**DOI:** 10.64898/2026.07.16.739011

**Authors:** Kai S. Shimagaki, John P. Barton

**Affiliations:** Department of Computational and Systems Biology, University of Pittsburgh School of Medicine, USA; Department of Physics and Astronomy, University of Pittsburgh, USA; Michigan Center for Applied and Interdisciplinary Mathematics, College of Literature, Science, and the Arts, University of Michigan, Michigan, USA

## Abstract

Epistasis, the non-additive effects of mutations, shapes fitness landscapes and evolutionary trajectories. Temporal genetic data reveal evolutionary dynamics and could be used to infer epistatic interactions, especially through linkage disequilibrium (LD) between interacting mutations. However, other evolutionary forces can also generate LD, challenging inference. Here, we systematically evaluated the accuracy of a variety of epistasis inference approaches across a range of selective pressures, recombination rates, and population sizes. In general, we found that inference accuracy depends on the evolutionary regime: methods based on marginal path likelihood (MPL) performed best under strong selection and low recombination, whereas quasi-linkage equilibrium (QLE) approaches were more accurate when recombination is frequent. We further showed that the strength of genetic drift can influence inference accuracy for approaches that learn from changes in allele frequencies over time. Collectively, our results show that the detectability of epistasis from temporal genetic data depends on the interplay between selection, recombination, and genetic drift, providing guidance for method selection across evolutionary contexts.

## Introduction

The concept of a fitness landscape, which maps genotype to reproductive capacity or fitness, is one of the most fundamental frameworks for understanding population evolution. This idea was introduced as an intuitive description of evolution as a hill-climbing process on a landscape, where regions of high and low survival correspond to peaks and valleys ^1,2^. Because fitness landscapes are inherently high-dimensional, where the dimension grows exponentially with sequence length, and because empirical information is limited, their practical use was historically restricted. However, recent experimental and theoretical advances have made quantitative estimation of fitness landscapes increasingly feasible, and their role in understanding evolutionary dynamics has become ever more central.

Epistasis, broadly defined as the non-additive effect of mutations on phenotype, including fitness, is central to understanding genotype-phenotype maps and fitness landscapes ^1,3,4^. When the fitness effect of a mutation depends on the surrounding genetic background, evolutionary trajectories become historically contingent. That is, because mutations that are beneficial in one genetic background may be neutral or deleterious in another, early mutations can constrain the set of accessible evolutionary paths. Such effects have been observed experimentally in diverse systems, including protein evolution and antimicrobial resistance ^5–14^.

Epistasis can be specifically defined in several ways, and different communities emphasize different aspects of the phenomenon ^15–17^. Throughout this work, we focus on specific (or idiosyncratic) epistasis, in which the fitness effect of a mutation depends directly on the allelic state of one or more other loci and can therefore be represented by explicit interaction terms in the fitness function. Other important forms of epistasis, including global epistasis arising from nonlinear genotype–phenotype relationships or higher-order interactions among multiple loci ^4,18–22^, are not considered here. Our goal is therefore not to distinguish among these different forms of epistasis, but rather to determine how accurately pairwise epistatic interactions can be inferred from evolutionary data under different conditions.

Practically, accurately estimating epistasis, a key determinant of fitness, can advance a wide range of fields ^3^. A deeper understanding of epistasis provides insight into historical contingency ^6,9–14^, and may help to predict how pathogens such as viruses evolve and escape from immune responses ^23–27^, as well as how drug resistant variants emerge ^28–30^. An accurate characterization of fitness and epistasis is also essential for guiding artificial evolution, including protein engineering aimed at developing proteins with desired functions ^31–33^.

Recent experimental and theoretical advances have made it increasingly possible to characterize epistasis and the structure of fitness landscapes. One important direction for estimating epistasis, which is the primary focus of the present work, is the analysis of time-series genetic data, whether from laboratory evolution experiments ^12,34–36^, ancient DNA ^37–39^, or the evolution of pathogens within or between hosts ^40–44^. These data reflect selective pressure and have become increasingly accessible. If epistatic interactions shape evolutionary dynamics, their effects may be reflected, at least indirectly, in the temporal patterns of allele frequencies and genetic correlations observed in these data ^41,42^.

Here, we study the performance of computational approaches for inferring epistatic interactions from temporal genetic data. We focus in particular on evolution from initially clonal populations, a condition that is common in experimental evolution and pathogen transmission with narrow bottlenecks. Intuitively, epistasis affects the frequencies of interacting mutation pairs, and measuring their covariance can provide insight into the underlying epistasis. However, inferring epistasis from temporal data is challenging because correlations between mutations may also arise from shared evolutionary history. For example, a selected genotype can sweep through the population while carrying neutral or deleterious mutations, resulting in elevated pairwise correlations even in the absence of positive epistatic interactions. Recombination further complicates inference: although it can break down correlations generated by genetic linkage and thereby reduce false-positive signals, it can also disrupt genuinely epistatically linked mutations, weakening signatures of epistasis. As a result, the detectability of epistasis depends on the evolutionary regime, including the strength of selection, the rate of recombination, and the magnitude of genetic drift ^45^.

Recently, numerous computational methods have been proposed to estimate epistatic interactions or signatures of epistatic fitness ^21,40,45–57^. However, only a limited number of these approaches have been applied to temporal genetic data ^40,54–57^. It remains unclear which methods most accurately detect epistatic interactions and under which evolutionary regimes they perform best. Furthermore, prior studies have not systematically examined inference accuracy across broad evolutionary regimes considering a range of selection strengths, recombination rates, mutation rates, and population sizes ^40,54–57^. This gap limits our understanding of the advantages and limitations of these inference methods. Here, we address this gap by evaluating the accuracy of epistasis detection across diverse evolutionary regimes using published methods that infer epistatic interactions from temporal multilocus genetic data without requiring intensive computation^54–57.^

We found that the accuracy of epistasis detection depends on the underlying evolutionary regimes, specifically the relative contributions of selection, recombination, and genetic drift. However, the timing of peak detectability is relatively insensitive when using frameworks incorporating statistics integrated over generations. In addition, we characterized the evolutionary regimes and linked them to conditions under which inference accuracy is high, using several statistical metrics.

## Results

### Simulations of population evolution on an epistatic fitness landscape

To explore the factors underlying the detectability of epistasis, we simulated evolution in a population of *N* individuals, where each individual’s genome is represented by a haploid, linear biallelic sequence ***g*** ∈ {0, 1} *L* of *L* loci, corresponding to a single linear chromosome. The population is initially clonal with genotype ***g*** = (0, …, 0), which we refer to as the wild-type. This initially clonal population assumption approximates scenarios such as the transmission of pathogens through tight population bottlenecks ^60,61^ and common experimental evolution setups. In our simulations, mutation and recombination can drive the emergence of new genotypes and diversify the initially clonal population. Population dynamics follow the Wright-Fisher (WF) process, including the effects of genetic drift scaling as *O*(1*/N* ) due to finite population size along with with natural selection, mutation, and recombination. Here we consider a fitness function *F* (***g***) which includes both additive effects of individual mutations and pairwise epistasis:

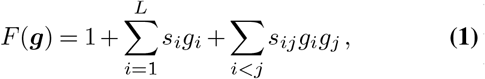

where the *s*_*i*_ and *s*_*ij*_ are selection coefficients and epistatic interactions within the same linear chromosome, respectively. Mutations can occur at any locus independently, each with the same mutation rate *µ*. Recombination can occur between any pair of genotypes ***g*** and ***g***^′^ at a crossover site *i*, with the uniform rate *r*, producing recombinant sequences of the form 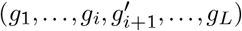.

Throughout this study, we set *L* = 50. We first consider a population size of *N* = 103 and a mutation rate of *µ* = 10−3. This focuses our initial investigation on a regime where the evolutionary forces of genetic drift, mutation, and selection are all expected to make comparable contributions to the dynamics. Later, we examine a broader range of population sizes and mutation rates. The magnitudes of additive and epistatic effects are characterized by a single parameter *σ*, such that | *s*_*i*_ | = | *s*_*ij*_ | = *σ*, which we define as the strength of selective pressure. We assume that the numbers of positive and negative coefficients are equal, and that epistatic interactions are sparse, with the number of nonzero elements scaling as *O*(*L*). Sequences are sampled every Δ*t* = 30 generations, and the corresponding time points are denoted by *t*_*k*_ = *k*Δ*t* where *k* ∈ {0, 1, …, *K*}. All simulations are run up to *t* = 10^3^ generations. Since genetic linkage, selective pressure, and recombination jointly influence the accuracy of epistasis inference, we vary the parameters *σ* and *r*. All statistics reported below are obtained from 30 independent simulations using the same underlying parameters, unless specified otherwise.

### Evolutionary regimes shape population structure

Before studying epistasis inference, we first quantitatively characterized how the underlying population structure changes under different evolutionary conditions. Adaptation from an initially clonal population has been studied extensively ^12,62–69^, revealing a transition from sweep-like evolution at low mutation supply to clonal interference when multiple beneficial mutations segregate simultaneously. Here, we quantify these evolutionary regimes using a set of complementary summary statistics describing allele-frequency dynamics, genetic diversity, fitness, and multilocus population structure. We then examine how these quantities depend on selection strength and recombination rate, providing a framework for interpreting the inference results presented below.

*Site-frequency classes* provide a simple summary of allele-frequency dynamics. We classify mutant alleles as highfrequency (*n*_high_*/L*) when their frequency exceeds 90%, low-frequency (*n*_low_*/L*) when it is below 10%, and polymorphic (*n*_poly_*/L*) otherwise. These classes show clear differences in simulations with weak and strong selection (**Fig. 1A-B**). Under weak selection, recurrent mutation primarily increases the number of polymorphic sites, while few mutations reach high frequency. In contrast, under strong selection, beneficial mutations and ones on high-fitness genetic backgrounds rise toward fixation, increasing *n*_high_ at the expense of polymorphic and low-frequency sites. Thus, weak selection maintains genetic diversity, whereas strong selection drives reduced diversity.

**Fig. 1.**
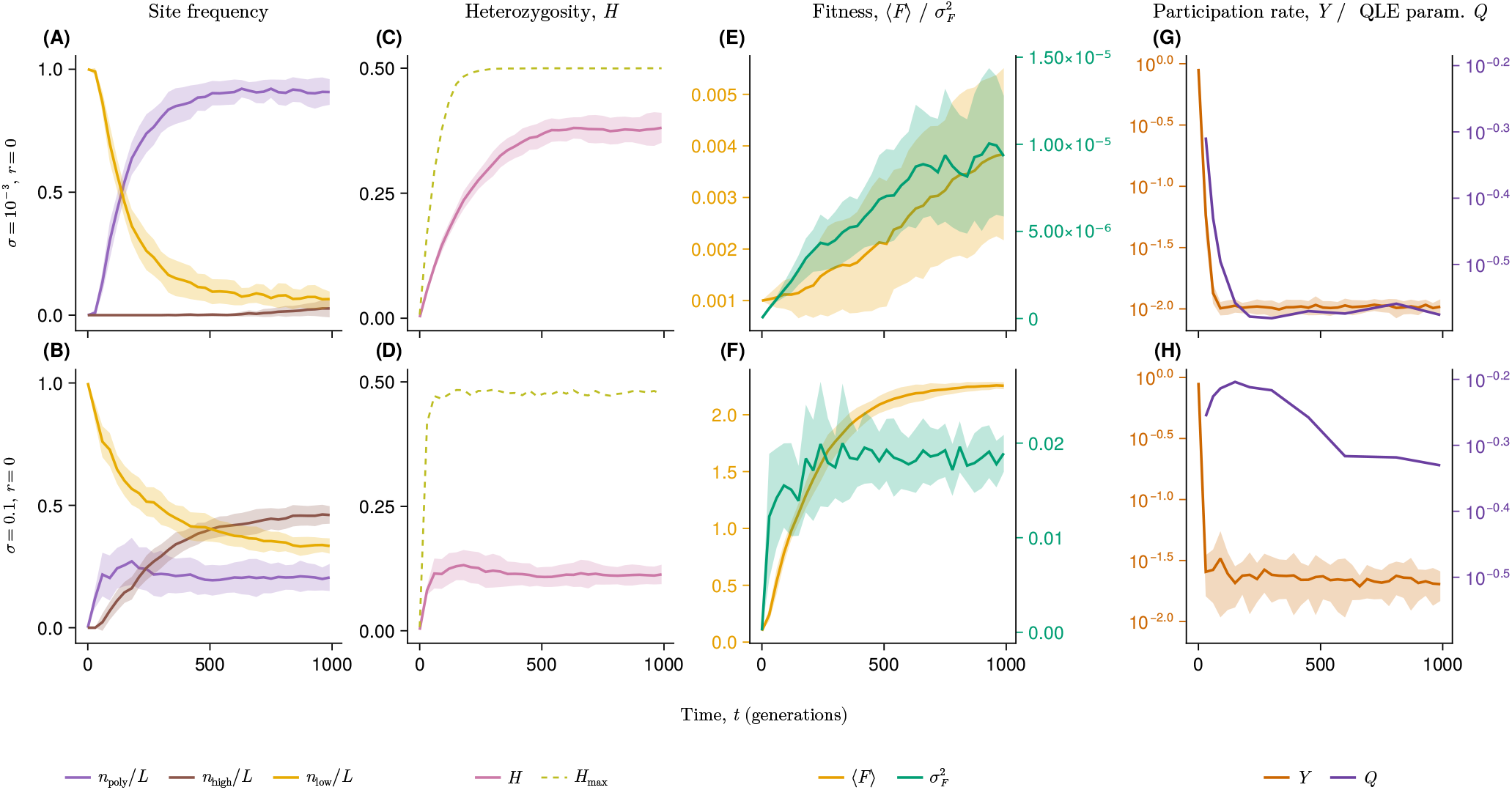
Dynamics of summary statistics for simulations with fixed population size *N* = 1000, mutation rate *µ* = 10^*−*3^, and recombination rate *r* = 0. Fractions of sites in each site-frequency class under weak (**A**) and strong (**B**) selection. Here, *n*_high_*/L* denotes the fraction of sites where the mutant allele frequency exceeds 90%, *n*_low_*/L* denotes the fraction below 10%, and *n*_poly_*/L* refers to sites with mutant allele frequencies between 10% and 90%. For all panels, lines show averages across 30 independent replicate simulations and shaded ribbons show standard deviations. (**C, D**) Heterozygosity under weak and strong selection, respectively. (**E, F**) Mean and variance of fitness under weak and strong selection, respectively. Because the fitness scale depends explicitly on *σ*, **E** and **F** use separate *y*-axis scales. (**G, H**) Multilocus statistics under weak and strong selection, respectively. Together, the participation ratio *Y* and QLE parameter *Q* quantify genotype-level concentration and allele-level correlations across loci.

*Heterozygosity, H* = ⟨ 2*x*_*i*_(1 − *x*_*i*_) ⟩ _*i*_ averaged across sites *i*, measures the average single-locus genetic diversity across the population. Under weak selection, mutation introduces new variation while alleles remain at intermediate frequencies, causing *H* to increase over time (**Fig. 1C-D**). Strong selection drives alleles toward fixation or loss, reducing the average heterozygosity despite the transient increase in diversity as sweeping mutations pass through intermediate frequencies. As a result, while individual loci may briefly approach the theoretical maximum *H*_max_ ≈ 0.5, the population-averaged heterozygosity remains substantially lower under strong selection.

*Mean fitness*, ⟨ *F* ⟩, and *fitness variance*, 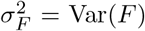, summarize adaptation at the genotype level (**Fig. 1E-F**). In a constant environment, the rate of adaptation is expected to be proportional to the genetic variance in population fitness according to Fisher’s fundamental theorem ^70,71^ (see **Supplementary Fig. S1** for an example comparison). As expected, we see that stronger selection produces higher mean fitness, even after normalizing by *σ*, indicating that this increase reflects more than just the scaling of the fitness function. In contrast, the normalized fitness variance, Var(*F* )*/σ*, is lower under strong selection because rapid fixation of high-fitness genotypes reduces standing genetic diversity, consistent with the decrease in heterozygosity (**Fig. 1C-D, Supplementary Fig. S2**).

*Participation ratio, Y*, and the *QLE parameter, Q*, are complementary statistics that characterize multilocus population structure. The participation ratio is given by

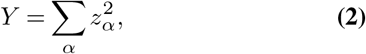

where *z*_*α*_ is the frequency of genotype *α* (ref. ^58^). This quantity measures genotype concentration (or “genetic condensation”) ^58^: maximally genetically diverse populations have *Y* ∼ *O*(1*/N* ), while populations dominated by a few genotypes have *Y* ∼ *O*(1). The QLE statistic,

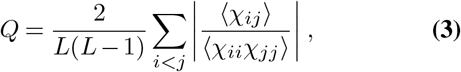

measures the average strength of pairwise associations among loci relative to single-locus diversity, where *χ*_*ij*_ = *x*_*ij*_ − *x*_*i*_*x*_*j*_ denotes the pairwise cumulant. Here the average ⟨·⟩ is taken over multiple simulations with the same underlying parameters. As discussed below, *Q* is closely related to the ratio *σ/*(*r* + 4*µ*) expected under quasi-linkage equilibrium (**Methods**), and therefore serves as a measure of departures from the QLE regime.

Although both statistics quantify population structure, they emphasize different aspects of the evolutionary dynamics. The participation ratio *Y* measures the concentration of genotypes, whereas *Q* measures nonrandom associations among loci. The two quantities therefore often vary together because the process of concentration on a small number of genotypes generates linkage disequilibrium. However, they are not necessarily equivalent. Selection and epistasis can produce substantial pairwise associations even in genetically diverse populations with relatively small *Y* .

As before, weak selection leads to genetic diversity, reducing both *Y* and *Q*. In contrast, strong selection amplifies high-fitness genetic backgrounds, increasing genotype concentration and strengthening multilocus associations (**Fig. 1G-H**, see also **Supplementary Fig. S2**). Unlike the single-locus statistics above, both *Y* and *Q* depend strongly on recombination, decreasing as recombination breaks down genetic associations (**Supplementary Figs. S3-S4**). Across a broader range of population sizes and mutation rates (**Supplementary Figs. S5-S7**), these statistics show that increasing mutation supply promotes genetic diversity, while stronger selection and weaker recombination favor genotype condensation and departures from QLE.

To summarize these trends across the evolutionary parameter space, we evaluated the summary statistics at later time points, after early transient dynamics (**Fig. 2, Supplementary Figs. S3-S4**). As expected, both *Q* and *Y* increase with stronger selection and decrease with increasing recombination, reflecting the transition from genetically diverse populations toward genotype condensation. In contrast, heterozygosity depends primarily on selection strength and is relatively insensitive to recombination. Together, these statistics show that selection and recombination produce distinct evolutionary regimes characterized by different patterns of genetic diversity and multilocus structure. In the following section, we study how these regimes influence the accuracy of epistasis inference.

**Fig. 2.**
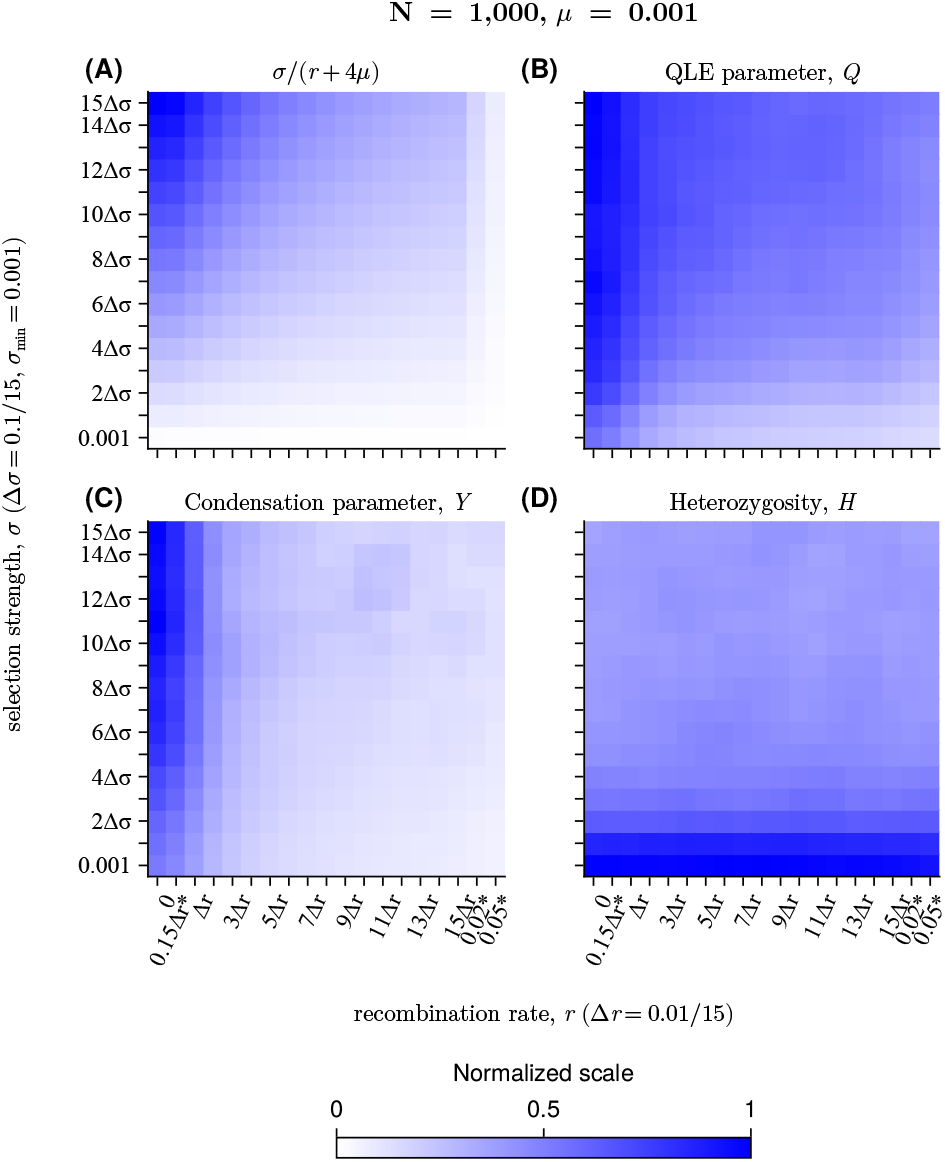
**(A)** The metric *σ/*(*r* + 4*µ*) increases when *σ* is larger and *r* is smaller. (**B**) The QLE parameter, *Q* defined as the sum over all pairs of the ratio of pairwise cumulants normalized by additive expectations, shows a similar dependence on *σ/*(*r* + 4*µ*). **(C)** The participation ratio, *Y* used as a measure of genotype condensation, also follows the same pattern. Its dependence on the recombination rate is sharp, with a rapid change in *Y* occurring when *r* between 5 × 10^*−*4^ and 5 × 10^*−*3^ (see also more quantitative analysis of genetic condensation as a function of recombination rate ^58,59^). **(D)** Heterozygosity, *H*, measures allele-level genetic diversity. Under strong selection and weak recombination, *Q* and *Y* values increase, meaning the system deviates from QLE, clonal interference becomes stronger, and genetic diversity decreases. The polarization of allele frequency and the decrease in diversity, measured by elevated *H*, occur as selection strength *σ* increases. All statistical values are evaluated at *t* = 10^3^ and averaged over 30 independent simulations at the fixed population size of *N* = 1, 000 and mutation rate *µ* = 10^*−*3^. For comparison across metrics, all values are normalized to range from 0 to 1. To smoothly estimate these statistics, we employed the 2D moving average method, where the value of the *i*th row and *j*th column is obtained from the raw values of neighboring *i*±1 rows and *j*±1 columns. Asterisk symbols * indicate elements that are not aligned on the linear scale of *σ* or *r* grids.

### Inference Frameworks for Epistasis Detection

To detect epistatic interactions from temporal genetic sequences, we considered four approaches: (1) a linkage disequilibrium (LD) statistic based on pairwise correlations; (2) a log-ratio of allele frequency method called universal footprint of epistasis (UFE) ^45,54^; (3) a quasi-linkage equilibrium (QLE) method derived from a steady state approximation; and (4) marginal path likelihood (MPL) method that explicitly models genotype frequency dynamics and linkage effects. All of these approaches make explicit use of the single-site and pairwise allele frequencies, *x*_*i*_(*t*) and *x*_*ij*_(*t*), where *t* indicates the generation at which the sequences are measured (see **Methods** for additional details).

#### Linkage Disequilibrium

One of the most straightforward approaches to quantifying non-random genetic association, including epistasis, is through the linkage disequilibrium metric (LD), which can be used to quantify epistatic effects with specific caution depending on the problems ^72,73^. The LD metric *D*_*ij*_(*t*) for interaction between loci *i* and *j* at generation *t* is given as

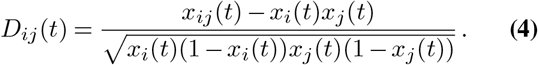

This metric is bounded between −1 and 1, making our analysis simpler. The version without normalization is also commonly used and referred to as the LD metric ^74^. Here, we used the normalized version to facilitate comparisons across mutation pairs that may have significantly different marginal allele frequencies.

#### Universal Footprint of Epistasis

To define the UFE metric, we first write the frequency of genotypes with alleles *a* and *b, a, b* ∈ {0, 1}, at focal sites *i* and *j* as *z*_*ab*_. The UFE metric is then given by ^45,54^

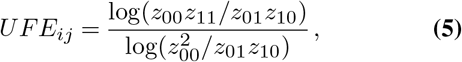

This metric is computed for all possible pairs *i* and *j*. This expression is also related to another widely used metric in genome-wide interaction analysis, log(*z*_00_*z*_11_*/z*_01_*z*_10_) (ref. ^46^).

#### Quasi-Linkage Equilibrium

QLE is a regime or “phase” in which LD rapidly converges to small but nonzero values, while single-locus allele frequencies evolve more slowly in time ^75^. This separation of time scales is expected when recombination and mutation act faster than the selective forces that drive allele-frequency change. QLE has been intensively investigated, mostly in two-locus systems, and has recently been generalized to multi-locus systems ^76,77^. In particular, multi-locus QLE has been reformulated under the assumption that the shape of the genotype frequency distribution is a Gibbs distribution with a pairwise energy function ^78^:

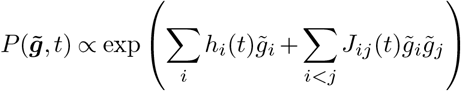

where *h*_*i*_(*t*) and *J*_*ij*_(*t*) are conjugate parameters of additive allele frequencies and pairwise cumulants. For consistency with prior literature, we represent WT/mutant alleles with 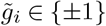 variables, rather than 0 or 1.

A series of equations involving fitness parameters, as well as *h* and *J*, naturally arise from the dynamics of the genotype distribution under selection, mutation, and recombination (Methods). Under the quasi-steady condition, where 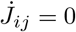 with small coupling | *J*_*ij*_ | ≪ 1, QLE yields the following relationship ^78^:

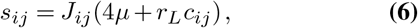

where *r*_*L*_ represents the probability that crossover occurs at any *L* − 1 links, *r*_*L*_ := *r*(*L* − 1), and *c*_*ij*_ represents the probability that alleles at the loci *i* and *j* are inherited from different parents ^78^ (see also its derivation Methods, as well as other models of recombination ^79,80^). The factor (4*µ* + *r*_*L*_*c*_*ij*_) represents the decay rate of LD values (ref. ^81^; see also Supplementary Information discussing LD dynamics for the two locus case.) Therefore, *J*_*ij*_ reflects a balance between epistatic forcing and the recombination/mutation driven relaxation.

The approach to inferring coupling interactions has been extensively studied in other biological problems with the developments of various frameworks ^82–85^. To estimate the coupling partners, we compute a covariance matrix 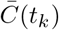 from the ensemble of sequences pooled over the time interval from *t*_0_ to *t*_*k*_, rather than by summing or integrating covariance matrices across time points ^79,80,86^. The coupling parameter is derived from the covariance within the mean-field approximation,

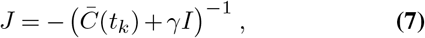

where *γ* controls the regularization strength. This form can be understood through a spectral decomposition of the covariance matrix, that is 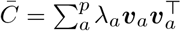 with eigenvalues *λ*_1_ ≥ · · · ≥ *λ*_*a*_ ≥ · · · ≥ 0 . Nonzero covariance arising from shared evolutionary history (e.g., genetic hitchhiking), often reflecting co-occurring mutations in the same background, contributes to the dominant part of the eigenspectrum (*λ*_*a*_ ≃ *λ*_1_), whereas more localized covariance signals, which may indicate co-evolutionary interactions (e.g., due to structural constraints in proteins), are associated with smaller eigen-modes (*λ*_*a*_ ≪ *λ*_1_). Therefore, inverting the covariance matrix can help suppress broad phylogenetic or linkage effects while enhancing the underlying epistatic signal ^87^. Based on Eq. (6), we expect that the QLE phase, characterized by small, steady LD and slowly changing additive frequencies, would be most appropriate when (4*µ* + *r*_*L*_*c*_*ij*_) ≫ |*s*_*i*_|, |*s*_*ij*_|.

#### Marginal Path Likelihood

QLE effectively assumes an infinite population size, as the genotype distribution does not fluctuate. However, when the population size is finite, genotype frequencies and their trajectories become stochastic. By denoting the genotype frequency vector as 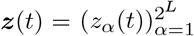, it is natural to consider the dynamics of *P* (***z***, *t*) under finite-population conditions. The principal idea of the MPL approach is therefore to infer fitness parameters by maximizing the likelihood of the observed genotype frequency trajectory. More formally, by denoting the entire genotype frequency trajectory, consisting of generations (*t*_1_, …, *t*_*K*_) as 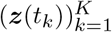, the MPL solution can be expressed as

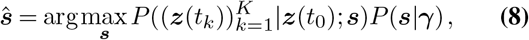

where ***s*** is a vectorized form of fitness parameters. *P* (***s*** | ***γ***) is a prior distribution, and ***γ*** is a hyperparameter, characterizing the uncertainty of parameters (see also Methods). The exact form of the path distribution is induced by WF process with selection, mutation, and recombination, in this study. Under the diffusion limit, where the mutation frequencies do not change abruptly within a short time interval Δ*t* (intuitively, *σ, µ, r* ≪ 1*/*Δ*t*), the trajectory distribution is approximated by a product of Gaussian terms. By leveraging the analytically amenable form of Gaussian products, the explicit expression of the MPL solution is obtained (Methods).

Finally, by switching coordinates from genotype frequencies, which live in an exponentially large space with respect to sequence length, to allele frequency space, using the relations *x*_*i*_ = Σ_*i*_ *z*_*g*_*g*_*i*_, *x*_*ij*_ = Σ_*i*_ *z*_*g*_*g*_*i*_*g*_*j*_ and so on (we consider *g*_*i*_ ∈ {0, 1}), the solution can be expressed as:

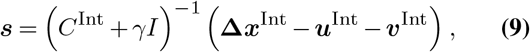

where *C* ^Int^ is the time-integrated covariance matrix, consisting of up to fourth-order terms, **Δ*x***^Int^ is the integrated allele frequency change, involving additive and pairwise terms over the trajectory, and ***u***^Int^, and ***v***^Int^ are the expected contributions from mutation and recombination, respectively. Unlike the QLE method, MPL can infer both additive and epistatic fitness components. Due to the robustness of the approach, it was applied to a broad range of data sets, including virus evolution within ^41,42,88^ and between ^43^ hosts, experimental evolution of bacteria ^89,90^, and high-throughput mutagenesis experiments ^91^.

### Patterns of linkage disequilibrium across time

We first asked when epistatic interactions become detectable during evolution. We began by considering the dynamics of LD, which can provide a signal about the presence or absence of epistasis between mutations in temporal data. Here, for simplicity we consider an asexually reproducing population (*r* = 0); results for the large-*r* case are reported in **Supplementary Fig. S8**.

Early in evolution, the distribution of LD values between mutations with positive or negative epistatic interactions, and between pairs of mutations that do not interact (“neutral” interactions) largely overlap. These distributions become more distinct at intermediate times, but further evolution causes them to blur together (**Fig. 3A-I**).

**Fig. 3.**
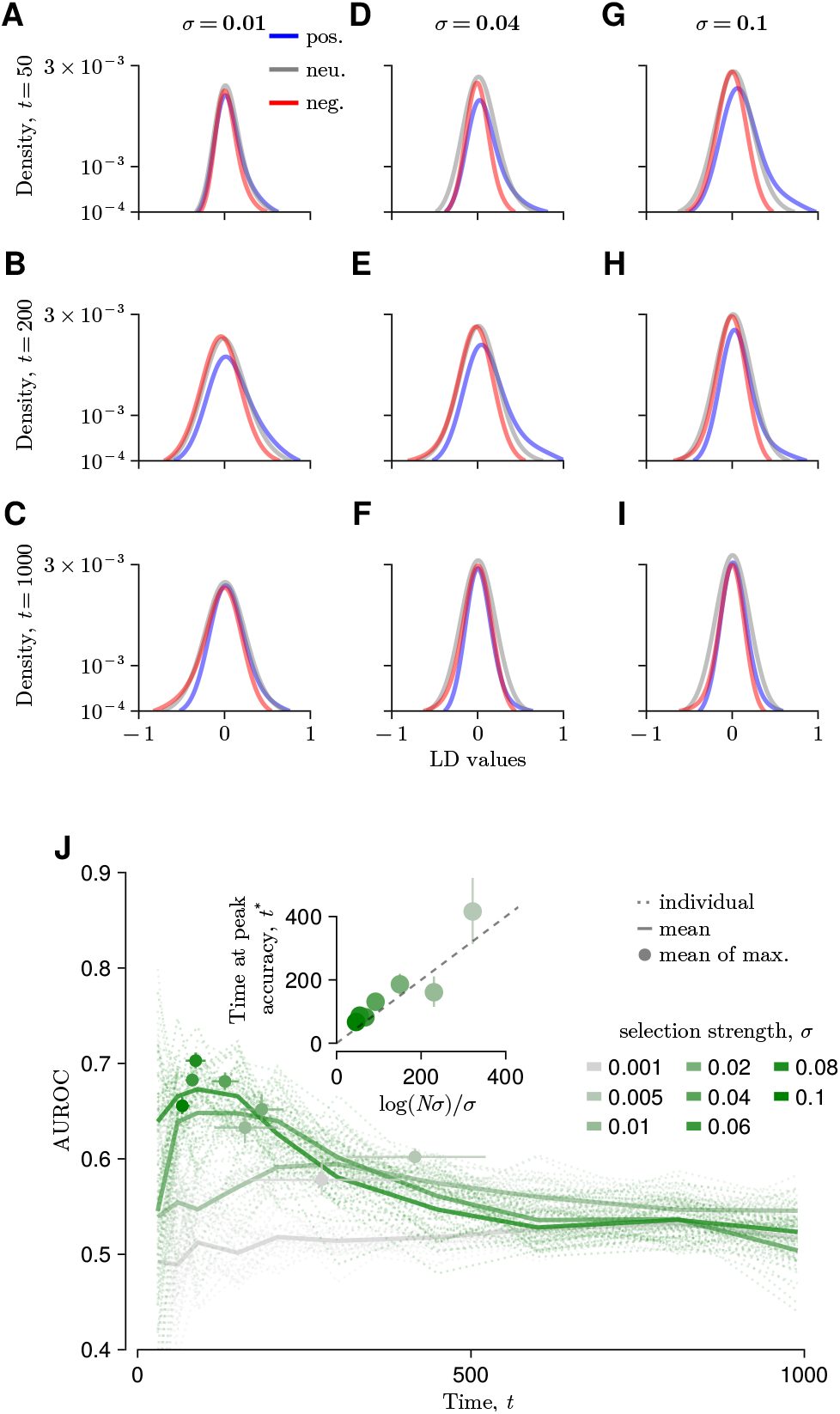
**A–I**, Distributions of LD values for pairs of mutations. The population size and recombination rate are *N* = 10^3^ and *r* = 0, respectively. The distributions tend to overlap during early generations. For weak selection (*σ* = 0.01; **A–C**), these distributions remain largely overlapping across a broad range of time points. Under intermediate selective pressure (*σ* = 0.04; **D–F**), the distribution associated with positive epistasis initially overlaps with the others but later becomes more distinct from the neutral and negative distributions. At longer times, the distributions begin to overlap again, though not as extensively as at the earliest stage. The case of strong selective pressure (*σ* = 0.1; **G–I**) shows a similar overall pattern to the intermediate case, but the period of strong deviation in the positive-epistasis distribution occurs earlier in the evolutionary timeline. **J**, AUROC values for detecting positive epistasis, used as an accuracy metric, tracked over time across different strengths of selection (*σ*). AUROC values tend to be higher at earlier times and gradually decrease along with *σ*. The timing of peak accuracy depends on the strength of selection: as selective strength increases, AUROC peaks become higher and occur earlier in evolution (inset). The same analysis for large *N* or frequent recombination is presented in **Fig. S11**. Elevated selection strength *σ* and population size *N* contribute to an earlier peak in accuracy (see also **Supplementary Tables 1-3**, which summarize additional quantitative tests based on the Kolmogorov-Smirnov test).

### Detection of epistasis via LD

Following the dynamics of LD described above, we found that the detection of positive epistasis based on LD was most accurate at earlier times, consistent with previous work ^45^ (**Fig. 3J**). This pattern is consistent across a broad range of selection strengths, particularly when selection dominates genetic drift (*σ* ≫ 1*/N* ).

More quantitatively, the accuracy of detecting positive epistasis via LD (*A*(*t*), quantified here via AUROC; see Methods for details) was well-approximated by the functional form

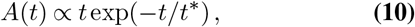

where the characteristic timescale coincides with the fixation time *t*_fix_ = log(*N σ*)*/σ* derived from a simplified model ^62^ (**Supplementary Fig. S9**). This phenomenological expression Eq. (10) provides the time at which the expected accuracy reaches its maximum, which is

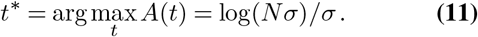

Intuitively, the first mutations that fix are likely to be driven by positive selection (including epistasis), and thus positively interacting pairs are likely to be enriched at this time. At later times, lineages with more complex patterns of positive and negative epistasis could rise to high frequencies, complicating the simple relationship between LD and positive epistasis (**Supplementary Fig. S10**). The estimated peak accuracy aligns with the numerical simulation for larger population sizes (**Supplementary Fig. S11**). However, this relationship no longer holds in the presence of recombination (**Supplementary Fig. S8**). Overall, the increase in selection strength pushes the time at which maximum separation of positive epistasis from neutral or deleterious epistasis to occur earlier, and a larger population size further enhances this trend (see also the analysis of the separation of LD distributions for positive, negative, and neutral epistasis in **Supplementary Tables 1-3**). The detection of negative epistasis via LD obeys displays a similar scaling in time, though accurate inference is much more challenging (**Supplementary Figs. S12-S18**).

### Performance of epistasis inference methods varies across evolutionary regimes

Having characterized the evolutionary regimes that emerge under different combinations of selection and recombination, we next examined how these regimes influence the accuracy of epistasis inference. As described above, we compared three approaches: a linkage disequilibrium (LD)-based metric, universal footprint of epistasis (UFE), the quasi-linkage equilibrium (QLE) method, and the marginal path likelihood (MPL) framework. To assess performance, we inferred epistatic interactions from simulated temporal genetic data and evaluated accuracy using AUROC for detecting epistasis (see **Methods)**. We first examined how accuracy evolves over time for different methods (**Fig. 4**).

**Fig. 4.**
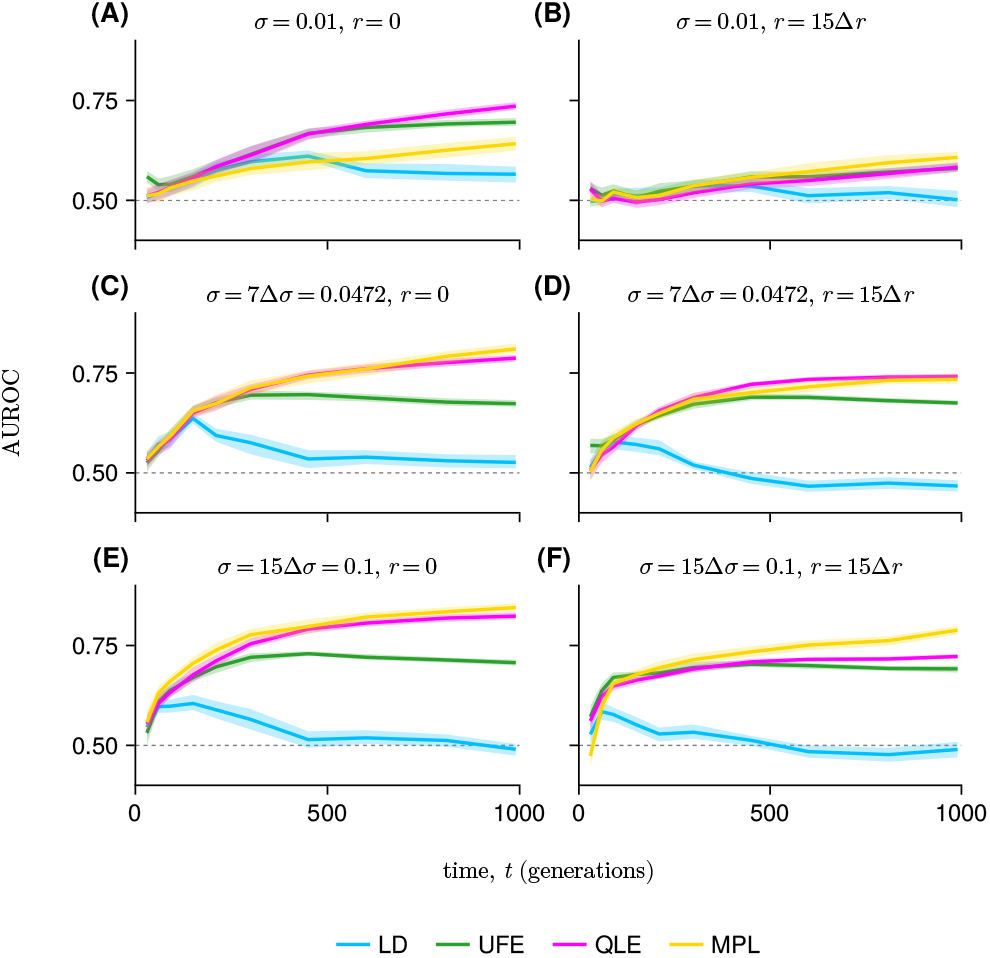
Epistasis inference accuracy over time for LD, UFE, QLE, and MPL at *N* = 10^3^, *µ* = 10^*−*3^. From top to bottom, panels show AUROC values for increasing selection strengths in the absence of recombination (**A, C, E**) and with *r* = 0.01 (**B, D, F**). Solid lines and shaded ribbons show the mean and standard deviation across replicates. The LD-based method performs well only within a narrow early time window, whereas UFE, QLE, and MPL maintain or improve accuracy over a broader range of time points. Overall, inference accuracy is lowest for the weakest-selection, high-recombination regime, where the epistatic signal is weakest and recombination reduces associations among loci.

At early times, the LD-based approach exhibits relatively high accuracy in detecting positive epistasis across a range of selection strengths and recombination rates. However, as background mutations accumulate and genetic linkage increases, detection accuracy declines substantially, consistent with the transient window identified above. In contrast, the detection accuracy for the QLE and MPL methods remains stable or increases as longer temporal windows are used (**Fig. 4, Supplementary Fig. S12**). These differences reflect the ways in which the methods use temporal information: LD relies on instantaneous correlations, whereas QLE and MPL incorporate information accumulated over time.

We next examined inference accuracy across a broad range of evolutionary parameters by evaluating performance at later times (*t* = 103) for different combinations of selection strength *σ* and recombination rate *r* (**Fig. 5**). Overall, the LD-based metric performs consistently worse than the other methods across most parameter regimes. In general, QLE and MPL achieve high accuracy across broad regions of parameter space, with particularly strong performance at higher selection strengths, whereas an increase in recombination generally reduces performance (see also the systematic effect of frequent recombination on inference accuracy in **Supplementary Figs. S15-S16**).

**Fig. 5.**
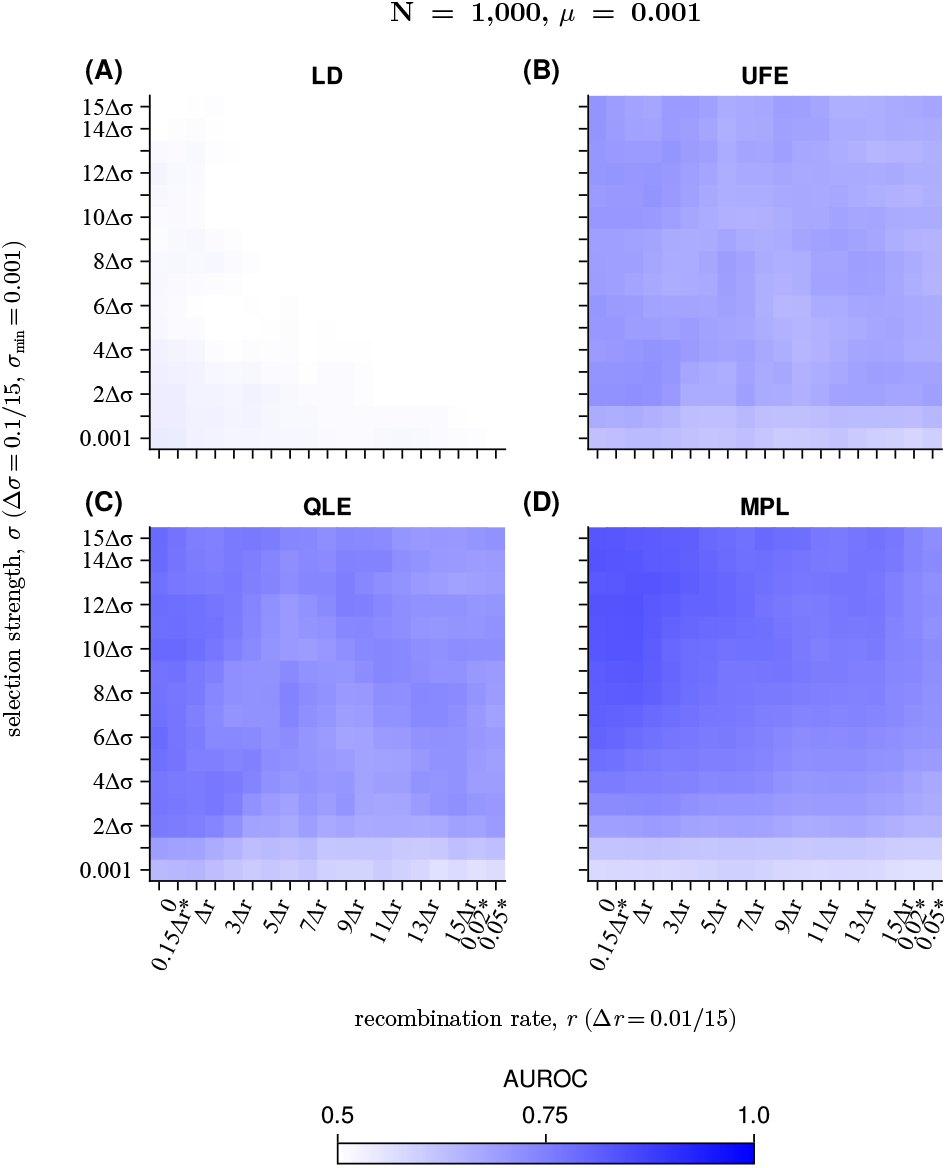
The AUROC values at a later evolutionary stage (*t* = 10^3^) across a range of selection strengths and recombination frequencies. Overall, the accuracy of QLE and MPL (**D**) remains high across broad parameter ranges, except for small *σ* and high *r*, with particularly better performance at higher selection strengths. As in **Fig. 2**, we applied a 2D moving average to smoothly estimate patterns of accuracy across parameter regimes.

We observed similar behavior for negative epistatic interactions (**Supplementary Figs. S12, S17-S18**). Across all methods, the dependence of inference accuracy on selection strength, recombination rate, mutation rate, population size, and sampling time is similar to that observed for positive epistasis. However, the absolute accuracy is slightly lower, particularly for the LD-based approach. One likely explanation is that, because our simulations begin from a clonal population, negatively interacting double mutants are less likely to accumulate to sufficiently high frequencies to generate a reliable LD signal.

The relative performance of QLE and MPL follows a consistent pattern across evolutionary regimes, which can be interpreted using the statistics introduced above. In regimes where the QLE parameter *Q* is large (indicating substantial linkage disequilibrium) and the participation ratio *Y* is elevated (reflecting reduced genetic diversity due to clonal interference), MPL generally achieves higher accuracy (**Fig. 6A**).

**Fig. 6.**
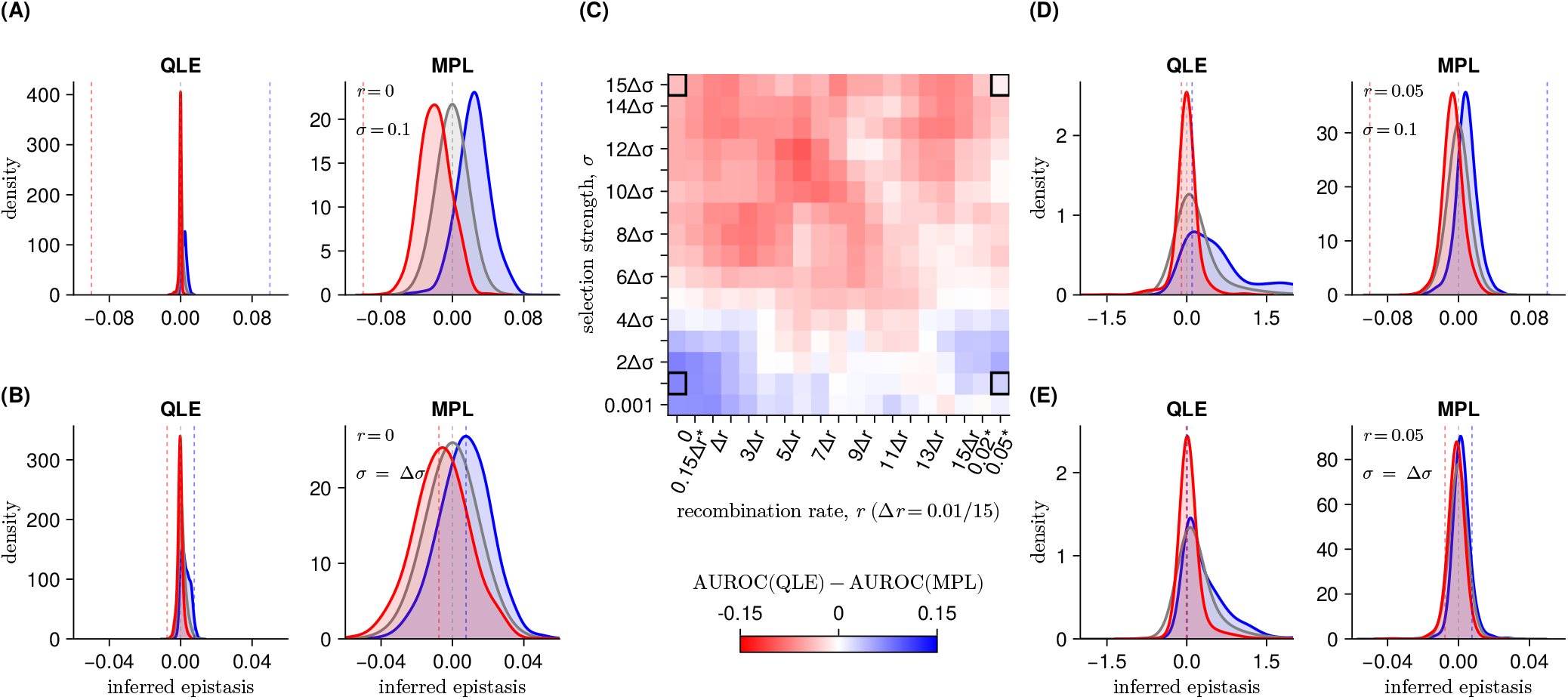
**(C)** Difference in AUROC values between QLE and MPL across parameter regimes. **(A-B, D-E)** Distributions of epistatic interactions inferred from sequences generated from 30 independent WF processes, using QLE and MPL frameworks. Dashed vertical lines represent the true underlying epistatic values. **(A) Large** *σ*, **small** *r*: Detection accuracy is highest for MPL in this regime, where strong selection and limited recombination create clonal interference. The performance of QLE improves relative to MPL in the regimes of **(B)** small *σ* and small *r* and **(D)** large *σ* and large *r*. **(E) Small** *σ*, **large** *r*: This regime is generally difficult for detecting epistasis, as frequent recombination breaks down most LD, including the LD generated by epistasis. Regularization parameters for both QLE and MPL are set to 1 throughout.

In these regimes, with large *σ* and small *r*, correlations between loci arise from linked evolutionary dynamics across expanding lineages, which MPL models explicitly. On the other hand, the performance of QLE is relatively stronger when *r* is smaller than, but roughly comparable to, the strength of selection *σ*, where both *Q* and *Y* attain moderate values (**Fig. 6A-B, D-E**). Both approaches struggle in the regime with small *σ* and large *r* (**Fig. 6E**). Here, accurate detection of epistasis becomes challenging as frequent recombination disrupts relatively weak signals of epistasis.

We found the relative performance of MPL and QLE in the regime with small *σ* and small *r* to be surprising. In prior work, MPL has been based on the assumption that selection is weak and the rate of recombination is low ^41,55,57^. However, in this regime QLE performed better at inferring underlying epistatic interactions, despite moderate values of *Q*. To understand this discrepancy, we next examined the role of genetic drift in shaping inference accuracy.

### Genetic drift can limit inference from allele frequency trajectories

One of the key differences between the MPL and QLE frameworks is the way that they use temporal information. The MPL approach infers selection coefficients and epistatic interactions from changes in allele frequencies over time. In contrast, QLE estimates epistatic interactions from covariance matrices that are averaged over multiple time points. As a result, QLE implicitly integrates information across time, whereas MPL depends more directly on short-term allele frequency changes.

Therefore, we hypothesized that noise due to genetic drift might impair the accuracy of MPL when the population size is small. Specifically, fluctuations in allele frequencies due to drift may obscure contributions of selection and epistasis, especially when the strength of selection is weak. By integrating information over time, QLE may be more insulated from the effects of drift.

To test this hypothesis, we systematically varied the population size *N* while holding the remaining evolutionary parameters fixed. We also tested comprehensive variation in the mutation rate *µ* to test the effects of mutation supply on inference. In larger *N* settings, the accuracy of MPL improved significantly (**Fig. 7, Supplementary Figs. S13-S18**). In particular, in the weak-*σ*, weak-*r* regime, MPL showed substantially better performance, indicating that the lower accuracy observed at *N* = 10 was largely due to genetic drift obscuring epistatic signals. This trend continues to smaller population sizes, where MPL performance declines. Like MPL, QLE improved accuracy with increasing *N* . However, the performance of QLE changed more modestly as a function of population size, reflecting its reliance on time-averaged statistics that are less sensitive to stochastic fluctuations. Comparatively, QLE tended to perform better at small population sizes, while MPL performed better at large population sizes, with a crossover around *N* ∼ 500 − 1000 depending on other evolutionary parameters. Both methods yielded more accurate inferences in populations with higher mutation rates.

**Fig. 7.**
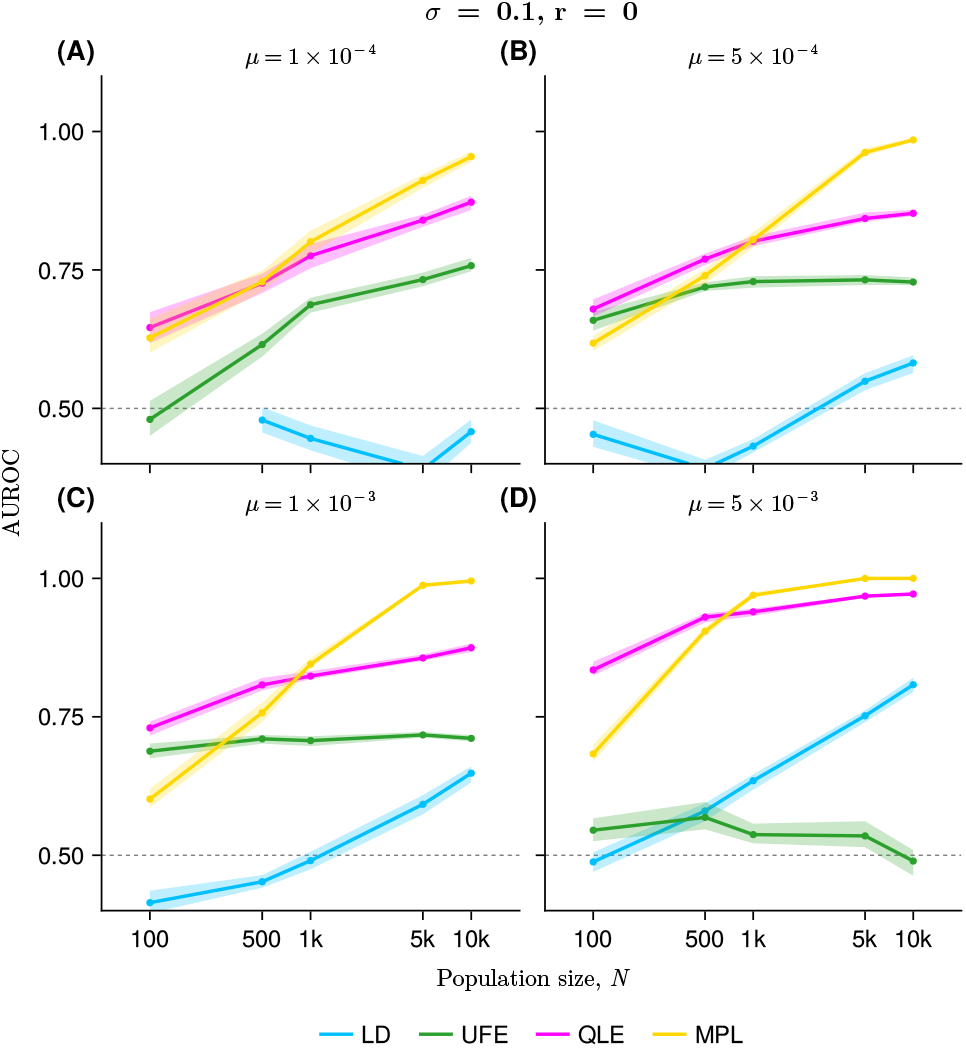
Effect of population size and mutation rate on epistasis inference accuracy under strong selection without recombination. AUROC values versus population size *N* at *µ* = 10^*−*4^ (**A**), *µ* = 5 × 10^*−*4^ (**B**), *µ* = 10^*−*3^ (**C**), and *µ* = 5 × 10^*−*3^ (**D**). Lines show the mean AUROC across 30 independent replicates, and shaded ribbons show the standard deviation. Inference accuracy generally increases along with population size. Typically, QLE is most accurate at small population sizes and MPL at large population sizes. At intermediate *N*, relative performance depends on other factors including mutation rate, selection strength, and recombination rate. The time dependence of the accuracy across different population sizes and mutation rates is shown in **Supplementary Fig. S14**.

Interestingly, the performance of UFE and LD as a function of population size was strongly modulated by the mutation rate. At the smallest mutation rate, UFE accuracy increased substantially along with population size. However, at high mutation rates, its performance actually declines as *N* increases. This may be due to the large number of mutation pairs with small but nonzero frequencies when *µN* is large. Because UFE uses the log-ratio of allele frequencies to infer epistasis, this approach may be especially susceptible to fluctuations in low frequency mutations. We observe the opposite trend for epistasis inference via LD: at low mutation rates, increasing *N* has little effect on accuracy, but at high mutation rates, accuracy increases substantially along with *N* . This may be linked to the difficulty in distinguishing LD generated by epistasis from chance associations, a challenge that the other three approaches are designed to overcome, when the mutation supply is low.

These results show that population size and mutation rate influence epistasis inference in different ways. Larger populations generally improve inference by suppressing genetic drift, although the magnitude and even direction of this effect depend on the statistical framework used. We next examine how these conclusions extend to the statistical power of epistasis detection.

### Statistical power for epistasis detection

To determine whether our conclusions depend on the choice of performance metric, we complemented the AUROC analysis with the true-positive rate at a fixed false-positive rate of 5% (TPR@5% FPR), which assesses statistical power. While AUROC summarizes performance across all decision thresholds, TPR@5% FPR quantifies the fraction of true epistatic interactions recovered while controlling the false-positive rate. This metric therefore provides a more stringent measure of detection power.

The TPR@5% FPR analysis confirmed the main results of our previous AUROC analyses (**Supplementary Figs. S21-S22**). For both positive and negative epistasis, inference accuracy generally improved with increasing population size, with MPL showing the largest gains as genetic drift was reduced. QLE performed well in smaller populations, although its performance improved more gradually with increasing *N* . Recombination reduced detection power across methods, and stronger selection generally increased it. Consistent with the AUROC analyses, positive epistatic interactions were detected more accurately than negative interactions, but with a similar relative dependence on selection strength, recombination rate, population size, and mutation rate.

### Inference of additive selection

We next asked whether the same evolutionary regimes similarly affect inference of additive fitness. Among the methods that we consider, only MPL directly estimates both additive and epistatic fitness parameters. We therefore compared MPL with a single-locus (SL) estimator that ignores correlations between loci ^41^,

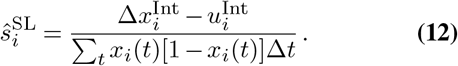

While QLE includes single-site fields *h*_*i*_, these fields are not directly equivalent to the additive selection coefficients *s*_*i*_ in the static inverse-Ising implementation used here ^92^. In classical QLE theory, additive effects enter through the dynamics of the single-site fields or allele frequencies and can include background-averaged contributions from epistatic interactions ^93^. We have therefore used QLE only to estimate pairwise epistatic coefficients.

Overall, additive selection was inferred substantially more accurately than pairwise epistasis across all parameter regimes (**Supplementary Figs. S19-S20**). This is expected because epistasis inference relies upon higher-order genetic correlations than for additive selection, which are correspondingly more difficult to resolve. As in the epistasis analysis, inference accuracy increased with selection strength and population size, while recombination had comparatively little effect. MPL consistently outperformed the SL estimator, particularly in larger populations where reduced genetic drift yields more informative allele frequency trajectories. These results indicate that accounting for multilocus genetic backgrounds improves inference even for additive selection, although recovering additive effects is generally less challenging than inferring epistatic interactions.

## Discussion

Temporal genetic data encode selective pressures, including features of the fitness landscape and epistatic interactions, yet inferring these effects remains challenging. Linkage disequilibrium (LD) provides a useful signal of both selection and epistasis, but correlations between mutations can arise from multiple sources, including genetic linkage and shared evolutionary history. As a result, distinguishing true epistatic interactions from background correlations depends sensitively on the evolutionary dynamics of the population.

Here, we tested four approaches for inferring epistatic interactions from temporal genetic data: linkage disequilibrium (LD), the universal footprint of epistasis (UFE) ^45,54^, quasi-linkage equilibrium (QLE) ^79,80^, and marginal path likelihood (MPL) ^41,55,57^. Overall, MPL and QLE consistently achieved the highest accuracy across the evolutionary regimes considered here, although no single method was uniformly optimal. Instead, inference accuracy depended on both the evolutionary regime and the statistical information used by each method.

These differences can be understood by considering the assumptions underlying each approach. LD and UFE rely primarily on pairwise genetic associations and therefore perform best when epistatic interactions generate detectable correlations that are not overwhelmed by background linkage or sampling noise. QLE infers epistatic interactions from covariance structure averaged across multiple time points, making it comparatively robust to stochastic fluctuations in allele-frequency trajectories. In contrast, MPL explicitly models temporal changes in allele frequencies and therefore benefits from large populations, where genetic drift is weak and evolutionary trajectories more faithfully reflect selection. Consequently, MPL performs best under conditions of strong selection, limited recombination, and sufficiently large populations, whereas QLE remains competitive in smaller populations and recombining populations that more closely satisfy its underlying assumptions.

Genetic drift emerged as one of the key factors determining inference accuracy. Trajectory-based methods such as MPL infer selection and epistatic effects from temporal changes in allele frequencies, and can therefore be sensitive to their fluctuations. When the population size is small, fluctuations due to drift can obscure contributions of selection and epistasis, reducing the accuracy of MPL even when linkage is strong. In contrast, methods based on covariance averaged across time, such as QLE, are less sensitive to these fluctuations. Increasing population size therefore suppresses drift and substantially improves MPL, while having a comparatively smaller effect on QLE.

Under our simulation conditions, we found that epistasis detection based on LD alone is limited to a transient temporal window early in evolution, in line with prior work ^45^. After a transient peak where LD distributions between epistatically interacting sites separate from one another, correlations between loci increasingly reflect random linkage rather than epistatic interactions, reducing detection accuracy. This behavior is well captured by a simple scaling heuristic, where the time of peak detectability coincides with the typical fixation time for selected mutations.

While we considered a range of inference methods and simulation conditions, our study also has several limitations. Our fitness model was rather simple, including only additive mutation effects and pairwise epistasis, while omitting higher-order interactions or global epistasis ^10,18,20,21,94–97^. This model also featured a fixed sparsity of epistatic interactions and uniform magnitudes of fitness effects. An increase in the density of true epistatic interactions increases the number of unknown parameters and strengthens the cumulative effects of interactions, likely making the inference of specific interacting sites more challenging. At the same time, sparse interaction networks result in a highly imbalanced inference problem, with few true interacting pairs among many possible ones. Systematically varying interaction density would therefore be a valuable extension of the present work. Our simulations used a clonal initial condition, one that may be well-suited to some biological scenarios. For example, tight transmission bottlenecks are common in the spread of pathogens, where one or a few individuals seed a new infection ^60^, and some experimental evolution studies use clonal starting populations. However, we also acknowledge that this condition is not universally applicable. An additional factor not considered here is the frequency with which genetic data are sampled over time. For methods such as MPL that rely on temporal changes in allele frequencies, the spacing between sampling time points can strongly influence inference accuracy ^41,55^. Finally, our analysis is based on simulated data, and further work will be needed to assess performance in empirical settings where these assumptions may not hold.

Our results also suggest several practical guidelines for experimental design. First, successful epistasis inference requires sufficient genetic variation. Larger effective population sizes and adequate mutation supply increase the number of informative polymorphisms and reduce the effects of genetic drift, improving the statistical power to resolve epistatic interactions. Second, epistatic signals must persist long enough to be observed. Extensive recombination can weaken informative associations between loci, implying that inference is more feasible in genomic regions or experimental systems where recombination is limited, or for interactions between more closely linked loci. Lastly, the choice of inference framework should reflect the underlying evolutionary regime. Trajectory-based approaches such as MPL are powerful but perform best when allele frequency changes can be measured accurately over time, while covariance-based approaches such as QLE remain effective in small populations and regimes with more limited temporal information.

## ACKNOWLEDGEMENTS

This work of K.S.S. and J.P.B. was supported by the National Institute of General Medical Sciences of the National Institutes of Health under Award Number R35GM161641. We thank two reviewers for their thoughtful suggestions. Computational resources were provided by the Computational and Systems Biology cluster at the University of Pittsburgh. The authors thank the system administrators for providing technical support.

## AUTHOR CONTRIBUTIONS

All authors contributed to the design of the study, interpretation of the results, and writing of the article. K.S.S. performed simulations and computational and mathematical analyses. J.P.B. supervised the project.

## Methods

### Modeling of population

We consider a haploid, bi-allelic population. Let 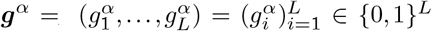 denote the genetic sequence of genotype *α*, consisting of *L* loci. The dynamics of population are characterized by selection, mutation, recombination, and genetic drift. Here, we define the fitness function that shapes the selective pressure and genetic dynamics.

#### Epistatic fitness function

Throughout this study, we assume the following pairwise fitness function:

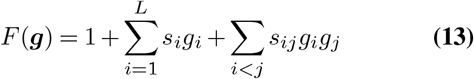

where *s*_*i*_ and *s*_*ij*_ represent additive selection coefficients and pairwise epistatic coefficients, respectively. For all nonzero coefficients, we set |*s*_*i*_| = |*s*_*ij*_| = *σ >* 0. The additive coefficients 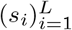 are drawn independently from a categorical distribution in which positive, negative, and zero values occur with equal probability (each with probability 1*/*3). Similarly, the pairwise epistatic coefficients (*s*_*ij*_)_*i<j*_ are drawn from a categorical distribution in which positive and negative values occur with probability 1*/L* each, and zero occurs with probability 1 − 2*/L*.

#### Wright-Fisher process

Given the fitness function, we model evolutionary dynamics using the Wright–Fisher (WF) process ^99,100^, a foundational framework in population genetics that describes how genotype frequencies change over time in a well-mixed population. The WF process is defined as a binomial or multinomial sampling process and naturally captures the effects of genetic drift. Let *z*_*α*_(*t*) denote the frequency of genotype *α* at time *t*. Denote by

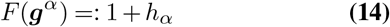

the fitness value *F* and genotype selection *h*_*α*_ of genotype *α*, and by *p*_*α*_(*z*) its effective survival probability under selection,

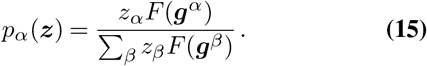

Then, WF process with the population size *N* can be expressed as the conditional probability of frequency *z* given frequency *z*^′^ at the previous time:

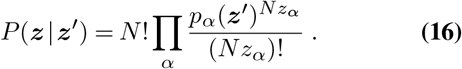

Due to the properties of the multinomial distribution, fluctuations in *z*_*α*_, quantified by the variance and covariance

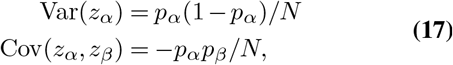

scale as *O*(1*/N* ). These fluctuations capture the effects of genetic drift. The changes in genotype frequency, denoted as 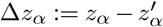, lead to the expected value over *P* (*z* | *z*^′^)

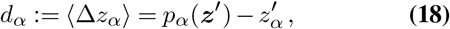

based on the properties of the multinomial distribution.

In the WF simulation, *p*_*α*_ is the survival probability after mutation, recombination, and selection. Each factor impacts the population in the following manner:

- Mutation generates new genotypes. Any genotype at any site can mutate uniformly randomly at a rate *µ* per generation per site across the population.
- Similarly, recombination creates new genotypes. In this case, we assume that any pair of genotypes can undergo recombination; they cross over genetic sequences at any site. For example, two genotypes 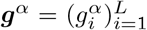 and 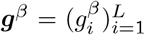 undergo recombination with a crossover point at *j*, then create new genotypes 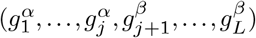, exchanging the genetic sequence of *α* and *β*.
- Lastly, selective pressure, characterized by fitness function, reshapes the distribution of genotypes. We randomly resample genotypes according to *p*_*α*_(*z*), the ratio characterized by the abundance of genotypes and fitness values.

In our simulations, all *N* individuals in the initial population start with wild-type sequences. The population then evolves over 1000 generations subject to selection, mutation, and recombination as described above.

### Quantify epistasis from temporal data

In this study, we used four metrics to measure the epistasis: (1) Linkage Disequilibrium (LD), (2) an approach called Universal Footprint of Epistasis (UFE), (3) a Quasi-Linkage Equilibrium (QLE)-based metric, and (4) a Marginal Path-Likelihood (MPL)-based metric. Below, we discuss the details of the UFE-, QLE-, and MPL-based metrics, which are relatively new and not commonly used to infer fitness parameters from populations. To describe these frameworks, we consider a bi-allelic, multi-locus population, which is realized under the WF process.

#### Universal Footprint of Epistasis

By encoding the wildtype and mutant alleles as 0 and 1, respectively, the mutation frequencies *z*_0_, *z*_1_, *z*_2_, and *z*_3_ correspond to the genotype frequencies of 00, 01, 10, and 11. The UFE metric is then defined as ^101,102^,

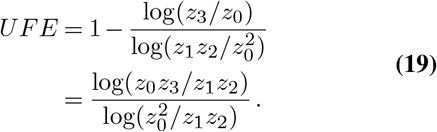

The numerator is equivalent to the metric used by Kimura ^103,104^, and the same metric has been used in recent genome-wide association study ^105^.

#### Quasi-linkage equilibrium

The QLE is a state where the LD decays rapidly and converges to smaller values (compare with additive frequencies), while the additive allele frequencies evolve relatively slowly, and these time-scales are distinctive. Under this QLE regime, the analytical calculations for population dynamics (i.e., the dynamics of genotype distribution) and for population-averaged statistics become simpler. QLE theory, which is based on an infinitely large population size, were intensively studied in population genetics. Recently, the multi-locus QLE theory was stimulated by mathematical analysis ^106^, using the statistical-physics formulation, which was subsequently applied to infer epistatic interactions by assuming Ising or Potts-type Gibbs distributions ^107–110^.

Here, we briefly summarize how epistatic interactions can be inferred within the QLE framework. The core concepts of the modern QLE theory are described in ref. ^106^, while its application to epistasis inference and detailed descriptions the QLE-based epistasis derivation can be found in refs. ^109,110^. A more general overview of QLE theory and its connections to MPL is provided in a recent study ^111^.

The general assumptions about genotype evolution, specifically how selection, mutation, and recombination act on the population, are the same as those introduced in the MPL section below. Here, however, we focus on large, well-mixed populations in which genetic drift is negligible and genetic linkage is weak, so correlations between loci are small. For consistency with previous QLE studies, we represent genotypes as ±1 sequences rather than sequences 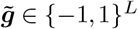. Under equilibrium conditions, the distribution of genotypes can be expressed in a Gibbs–Boltzmann–type form with a pairwise energy function given by:

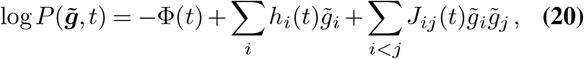

where Φ is the normalization constant of the distribution, and *h*_*i*_ and *J*_*ij*_ are the conjugate parameters associated with the allele frequency *x*_*i*_ and the pairwise term *x*_*ij*_ (or, equivalently, with its covariance *C*_*ij*_ = *x*_*ij*_ − *x*_*i*_*x*_*j*_). Here, *h*_*i*_ ∈ ℝ, ∀*i* ∈ {1, …, *L*} is distinct from the genotype level selection *h*_*α*_, ∀*α* ∈ {1, …, 2^*L*^}, which appears in the MPL framework. We shall refer to *h*_*i*_ as additive parameters and to *J* as coupling parameters. Empirically, this model can reproduce higher-order statistics in long-term protein evolution ^112,113^, and multiple computational frameworks were developed to infer these parameters from real genetic data ^114–121^ .

The epistatic coefficients can be inferred by estimating the coupling parameters *J* and using the relationship between the fitness model and the equilibrium distribution, as described below. To clarify this relationship, the dynamics of the genotype distribution and its steady state were considered. As in the forward WF simulations and the derivation of the MPL framework below, genotype dynamics are governed by selection, mutation, and recombination. Accordingly, the time evolution of the genotype probability distribution, 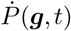, except that here we work with the probability of genotype *g* rather than its frequency *x*, and present the formulation more explicitly.

The central difference is that the QLE-based epistasis inference framework explicitly uses the genotype distribution *P* (*g, t*) to obtain its time derivative. The explicit expression of *P* (*g, t*) given Eq. (20) becomes

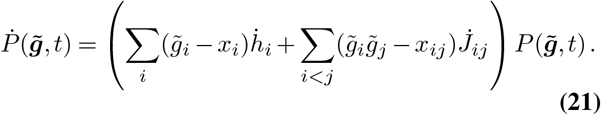

The equation Eq. (21) can be equated with the dynamics of *P* (*g, t*) based on the dynamical rule, defined by the selection, recombination, and mutation, for which we explicitly write the terms for the first and second terms.

Next, the dynamics of the genotype probability 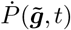 can be derived by considering each evolutionary mechanism: selection, mutation, and recombination. Much of the following discussion is based on ref. ^106^, while the contribution of mutation, which was not fully developed there, is incorporated following subsequent studies refs. ^109,110^.

First, the effects of selection. Consider that the fitness changes the population frequency exponentially, and after a short time period Δ*t*, the genotype distribution changes such that

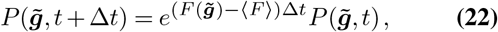

where 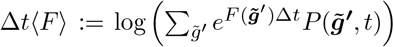. Within a short interval (|Δ*t*(*F ™* ⟨*F*⟩)|≪ 1) and to leading order in *O*(*F* Δ*t*), the change in the distribution due to selection is given by

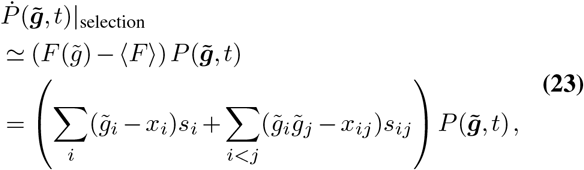

where ⟨*F*⟩ denotes the average of *F* over the probability 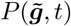.

Next, we consider mutation effects. As in the MPL framework below, under a low mutation rate, at most a single mutation occurs per genotype, driving changes in *P* (*g, t*) according to

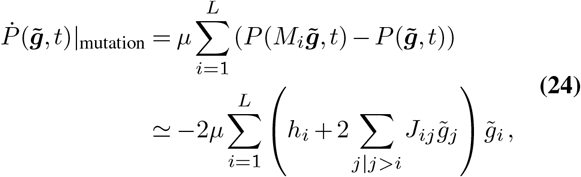

where *M*_*i*_ denotes an operator acting on 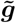 that flips the sign of the allele at site *i* such that 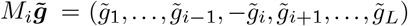. The final result uses the assumption that |*h*_*i*_| and |*J*_*ij*_| are much smaller than 1 for all ∀*i, j* .

Finally, we consider recombination effects. Let 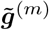, and 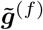 denote the genotypes inherited from the two parents. We also introduce an auxiliary variable *ξ* ∈ {0, 1}^*L*^, which specifies the parental origin of each allele: the allele at site *i* is inherited from parent *m* if *ξ*_*i*_ = 1 and from parent *f* if *ξ*_*i*_ = 0.

The offspring genotype 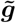 is then given by

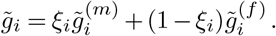

By denoting *C*(*ξ*) as a probability of crossover pattern *ξ*, the probability that loci *i* and *j* are inherited from different parents is given by ^106^

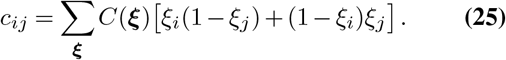

Under infrequent recombination, where crossover occurs only once per paired sequence, *c*_*ij*_ can be expressed as *c*_*ij*_ = |*j* − *i*| */*(*L* − 1) (ref. ^111^; see also ref. ^122^) . Although we consider the crossover of genetic sequences, previous studies also examined free recombination scenarios where swapping of alleles occurs locus-by-locus independently ^107,110^.

With these definitions, the contribution of recombination to the time evolution of the genotype probability distribution can be written as (for more detailed derivations, see Appendix C of Ref. ^106^ or Sec. 3 of Ref. ^109^.)

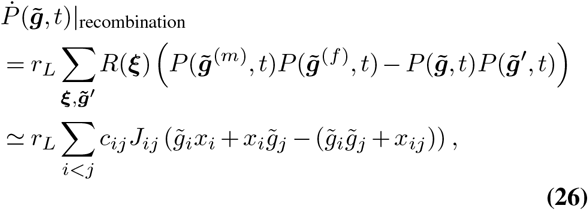

where *r*_*L*_ := *r*(*L* − 1) . By equating the direct time derivative of 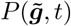 in Eq. (21) with the contributions from the evolutionary forces summarized in Eqs. (23–26), for arbitrary genotypes 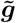, we obtain the following relationships (which are presented as Eqs. (24) and (25) in Ref. ^123^, and Eqs. (18) and (19) in Ref. ^109^):

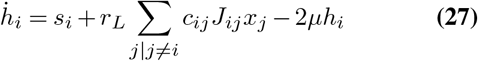

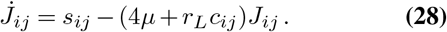

Therefore, under the steady state in *J*_*ij*_ we obtain the relation, which yields the expression shown in the main Eq. (6) and allows us to estimate *s*_*ij*_ via *J*_*ij*_:

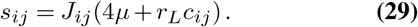

The statistical framework for inferring the coupling parameters *J* under the assumption of a steady-state distribution is well established and is known as the inverse Ising (or Potts) problem. In this study, we infer *J* from an ensemble of sequences using a simple mean-field approach:

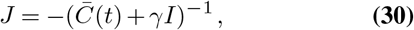

where the covariance matrix 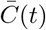 is obtained from the ensemble of sequences across multiple time points from the initial to the generation *t*, and *γ* is regularization parameter. In this study, we pool all time points together, following previous QLE-based epistasis inference works ^107,109,110^.

#### Marginal path likelihood

The MPL approach is derived from faithfully expressing the WF process under the diffusion approximation, where the frequency change is small over time, and maximizing the evolutionary parameter over the evolution. In the diffusion limit, we assume the evolutionary parameters are small, specifically *s*_*i*_, *s*_*ij*_, *µ* and *r* are of order (1*/N* ). Given the fitness function and WF process defined above, the diffusion approximation replaces the WF process with a diffusion process in which the transition probability *P* (*z*(*t*_*k*+1_) *z*(*t*_*k*_)) is modeled as a Gaussian distribution at time *t*_*k*+1_, conditional on the population state at time *t*_*k*_. Accordingly,

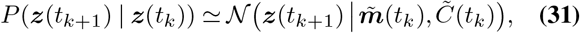

where 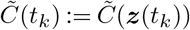 is the covariance matrix whose entries are given by the variances and covariances in Eq. (17), and 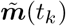 is the mean frequency vector at time *t*_*k*+1_, defined as

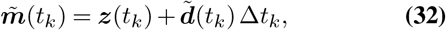

with Δ*t*_*k*_ := *t*_*k*+1_ − *t*_*k*_ and 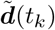 denoting the expected change in frequency, as given in Eq. (18).

Since the evolutionary parameters are small (order of *O*(1*/N* )), the covariance and mean vector can be further simplified by taking only the leading terms of order *O*(1*/N* ), which are given as

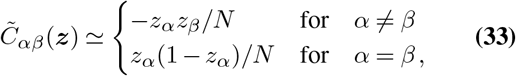

and

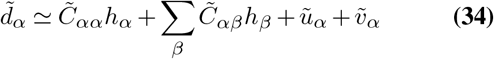

where 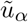 and 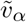 are expected changes in genotype frequency due to mutation and recombination, given as

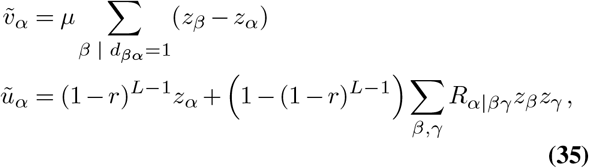

where *d*_*αβ*_ and *R*_*α*|*βγ*_ denote the Hamming distance between genotypes *α* and *β*, and the probability that recombination between genotypes *β* and *γ* produces genotype *α*, respectively . 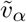 represents the frequency change due to mutations; genotypes *β* can mutate into *α*, while genotype *α* can also mutate into other genotypes. 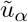 represents contributions from recombination: the first term corresponds to the absence of recombination (none of the *L* − 1 recombination events occur), and the second term corresponds to recombination between genotypes *β* and *γ* generating a new genotype *α*.

The optimal fitness parameters are then obtained by maximizing the full path-likelihood function under the diffusion limit, which can be derived analytically by exploiting the linearity of the dynamics in *s* and the Gaussian structure of the likelihood. Consequently, the optimal 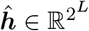 can be expressed analytically as:

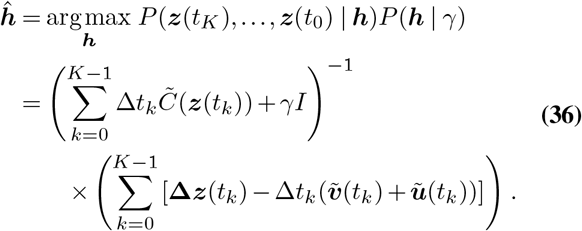

Here, *P* (***h*** | *γ*) denotes the prior distribution of the genotype selection parameter, which we take to be a normal distribution with zero mean and variance *γ*^−1^. The final result holds under the assumptions that the dynamics follow a WF process, the evolutionary parameters remain constant over time, and the genotype-frequency changes between successive time steps are sufficiently small for the diffusion approximation to remain valid.

To obtain the allele-level selection and epistatic coefficients *s*_*i*_ and *s*_*ij*_, we project the quantities defined in genotype space onto allele space. Let *G* be a binary matrix that maps additive mutations and mutation pairs to genotypes, such that

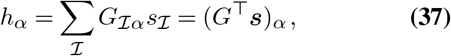

where index I runs over individual site (*i*) and pairs of sites *i* and *j*, and ***s*** = (*s*_1_, …, *s*_*L*_, *s*_1,2_, …, *s*_*L*−1,*L*_) . Given the *G* matrix, allele frequency ***x*** can be expressed as

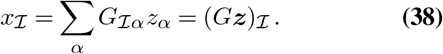

By denoting the covariance given as

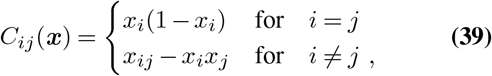

which is related to the genotype covariance matrix 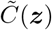 through the *G* matrix such that

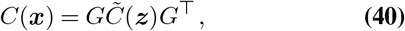

and using Eq. (37) as well as Eq. (38), we obtain the optimal selection in allele space. Multiplying the equation by the matrix *G* from the left yields the genotype selection in Eq. (36), that is, 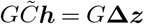.

Therefore, the optimal selection coefficient in allele level is obtained as

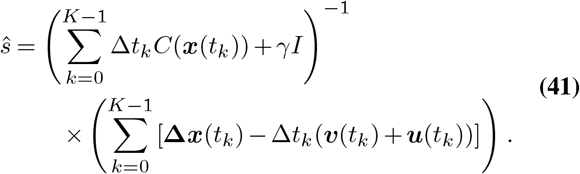

The frequency changes due to mutation and recombination, denoted as 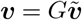 and ***u*** = *G****ũ***, are obtained by projecting the *G* matrix from the left. Integrating ***v*** and ***u*** yields ***v***^Int^ and ***u***^Int^ that are used in Eq. (9). The final result in Eq. (41) is computationally tractable even for long genetic sequences, and we use this equation to infer the epistatic coefficients. For additional technical details, please refer to the prior works ^122,124,125^.

### AUROC calculations

Here, we used AUROC as one quantitative measure of the accuracy of epistasis inference. Because the true epistatic coefficients take three possible values, *s*_*ij*_ ∈ {± *σ*, 0}, we evaluated positive and negative epistasis detection as separate one-vs-rest binary classification problems. For positive epistasis, pairs of mutations are sorted in descending order based on their corresponding inferred epistatic interactions. These mutations are then transformed into boolean values: true if the true epistatic interaction is positive (*s*_*ij*_ *>* 0), and false otherwise (i.e., one-vs-rest). Similarly, to obtain AUROC values of negative epistatic interactions, mutations are treated as true if corresponding true epistatic interactions are negative, and false otherwise. The AUROC values for selection coefficients are obtained in the same manner. If the inference is perfect, that means all true positives appear first, followed by all false positives, resulting in an AUROC of 1.0. In contrast, if true positives appear uniformly at random, the AUROC value approaches 0.5. The AUROC values are obtained from independent simulations, and the mean and standard deviation of the AUROC values are calculated across the 30 replicates. If pairwise frequency remains low over time or if pairs of mutations are never observed, inferring epistasis and analyzing the detectability of epistasis is impractical ^122^. Therefore, we excluded the pairs of mutations with an average pairwise frequency *x*_*ij*_ below 0.05 over time.

### QLE parameter

Here, we analyze the dynamics of linkage disequilibrium (LD) system, defined as *D*_*ij*_ = *x*_*ij*_ − *x*_*i*_*x*_*j*_, under the combined effects of mutation, recombination, and selection. For simplicity, we focus on a two-locus system (*D* = *x*_12_ − *x*_1_*x*_2_), noting that the results generalize to multilocus settings. We consider four genotypes, ‘00’, ‘01’, ‘10’, and ‘11’, where 1 denotes a mutation and 0 the wild-type. Let their genotype frequencies be indexed as (*z*_0_, *z*_1_, *z*_2_, *z*_3_), which by definition satisfies 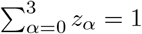. These genotype frequencies can be expressed in terms of the marginal allele frequencies *x*_1_, *x*_2_ and the pairwise allele frequency *x*_12_ as follows:

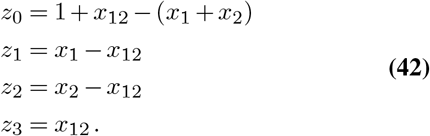

First, we consider the effect of recombination. The contribution of recombination to the genotype frequencies is given by (ref. ^104^):

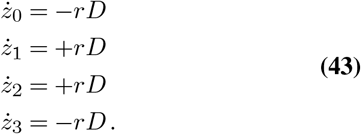

Because recombination alone does not change single-locus allele frequencies (i.e., 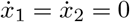) due to recombination, we obtain

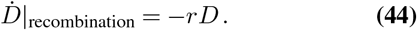

Second, we consider the effect of mutation. Assuming a uniform mutation rate *µ* across loci, the resulting changes in allele frequencies due to mutation are given by

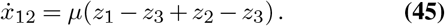

Here, the mutation terms describe how mutations in geno-types “01” and “10” increase the frequency of genotype “11”, while any mutation occurring in genotype “11” decreases its frequency. The resulting change in the single-locus allele frequencies is given by 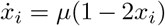. Accordingly, the change in LD due to mutation is given by

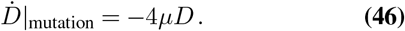

Finally, we consider the effects of selection. Selection changes genotype frequencies according to

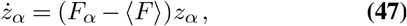

where ⟨*F*⟩ = Σ_*β*_ *F*_*β*_*z*_*β*_ is the mean fitness of the population. The resulting changes in the marginal and pairwise allele frequencies due to selection are given by: 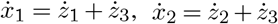, and 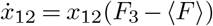. Accordingly, under the QLE assumption where *D* is small, the contribution of selection to the LD dynamics, 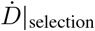, can be expressed as

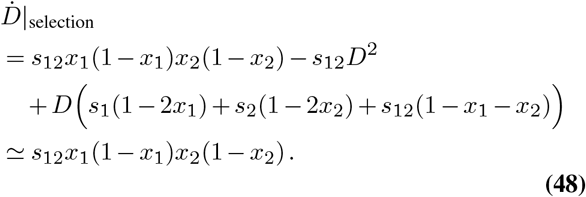

Combining all contributions yields the net change in *D*, given by

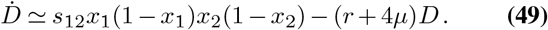

To make the connection with the main text explicit, we switch notation as follows: *D* → *χ*_*ij*_, *x*_*i*_(1 − *x*_*i*_) → *χ*_*ii*_, and *s*_*ij*_ → *σ*. With these substitutions, the relation can be written as

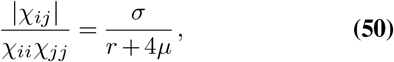

The left-hand side in Eq. (50), representing LD per unit heterozygosity, does not explicitly depend on model parameters, whereas the right-hand side depends only on the parameters *σ, r*, and *µ*. This equation therefore characterizes the typical behavior of the statistic *χ*_*ij*_*/*(*χ*_*ii*_*χ*_*jj*_) across parameter regimes. Moreover, it directly implies that when *σ* ≪ *r* + 4*µ*, the condition |*χ*_*ij*_ | ≪ *χ*_*ii*_*χ*_*jj*_ holds, and the population lies in the QLE regime. An equivalent relationship can be derived for multilocus systems within QLE theory using a Gaussian closure scheme ^111,126^.

## Code availability

Computer code used in this study is available in the GitHub repository of https://github.com/bartonlab/paper-epistasis-detectability.

## Supplementary Information

**Supplementary Fig. S1.**
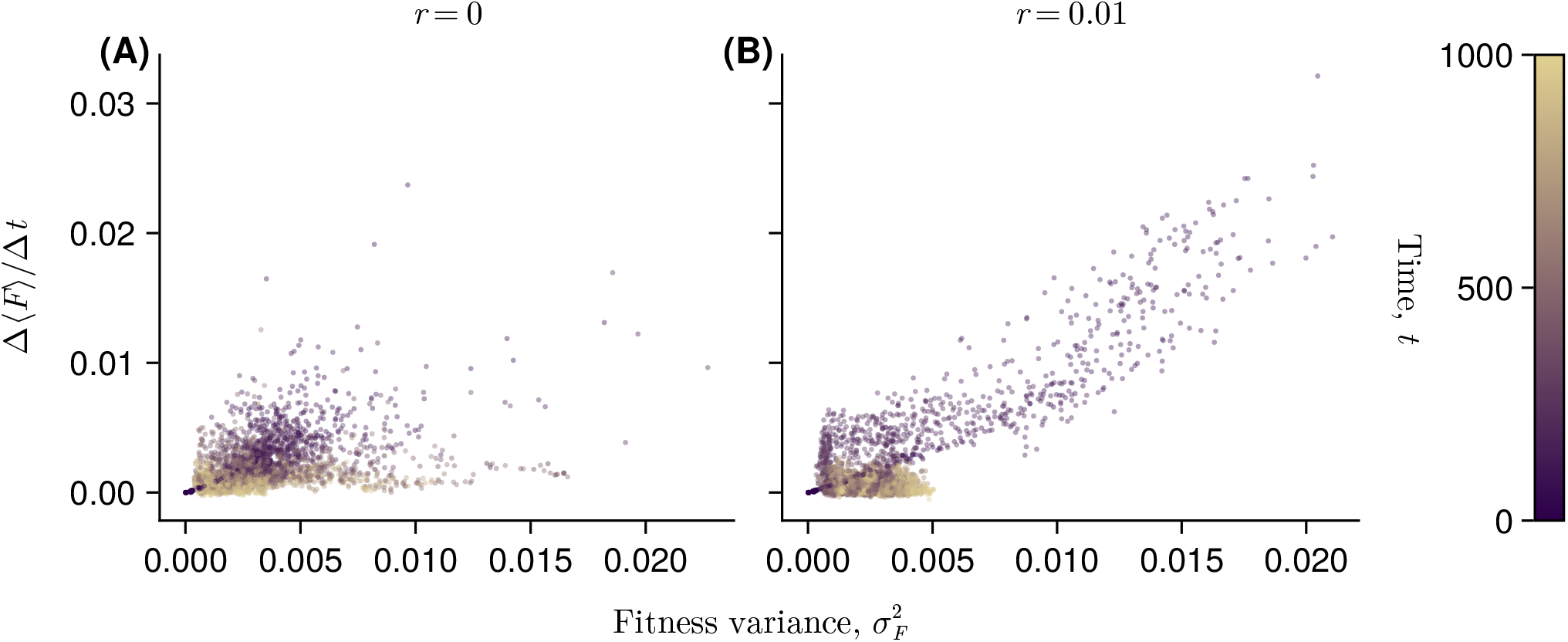
Shown are low-drift, strong-selection conditions (*µ* = 10^−4^, *N* = 10^4^), where selection is the dominant driver of changes in mean fitness ⟨*F* ⟩. The rate of mean fitness change, Δ⟨*F* ⟩*/*Δ*t*, is estimated from numerical changes over Δ*t* = 30 generations. Each point represents the result from one of the 30 replicates. Panel **A** shows the asexual case (*r* ), whereas panel **B** shows frequent recombination (*r* = 0.01), which leads to a rapid increase in ⟨*F* ⟩ and large 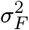. Overall, Δ⟨*F* ⟩*/*Δ*t* and 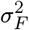 are approximately linearly correlated, particularly at early generations.

**Supplementary Fig. S2.**
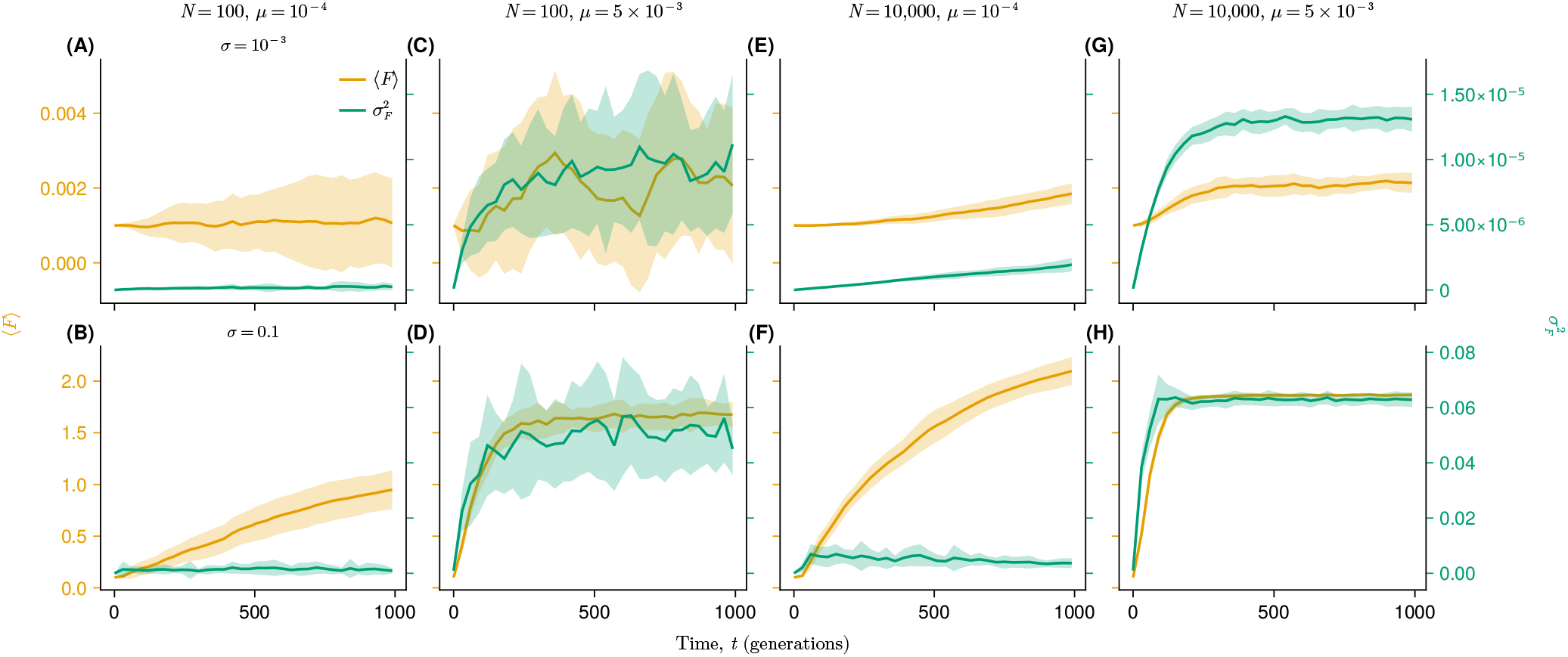
Same as above, these results are derived from asexual evolution (*r* = 0). Overall, smaller-*σ* cases (panels **A, C, E**, and **G**) maintain lower mean fitness and fitness variance, whereas larger-*σ* cases (panels **B, D, F**, and **H**) push the average fitness to higher values. This remains true even after normalizing the mean fitness by *σ*, i.e., using ⟨*F* ⟩*/σ*, to remove the explicit *σ*-dependence of the fitness function. For example, the ratio of mean fitness between the large- and small-*σ* cases is approximately 1.5*/*0.002 = 750, which is much larger than the ratio of the *σ* values themselves, 0.1*/*0.001 = 100. This suggests that larger *σ* not only rescales fitness, but also changes the population dynamics by more efficiently amplifying high-fitness genotypes. The normalized fitness variance, 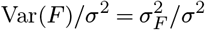, shows a more complex pattern. Strong selection can rapidly push genotypes toward fixation or extinction, thereby reducing standing diversity. Consequently, the normalized variance can be lower in the strong-selection case. For example, in a single-locus case, the fitness variance is proportional to *σ*^2^*x*(1 − *x*), which becomes small when the allele frequency *x* approaches either 0 or 1. This is consistent with the heterozygosity results: strong selection, especially at large *N*, can produce lower average heterozygosity *H* than weak selection because allele frequencies become more polarized.

**Supplementary Fig. S3.**
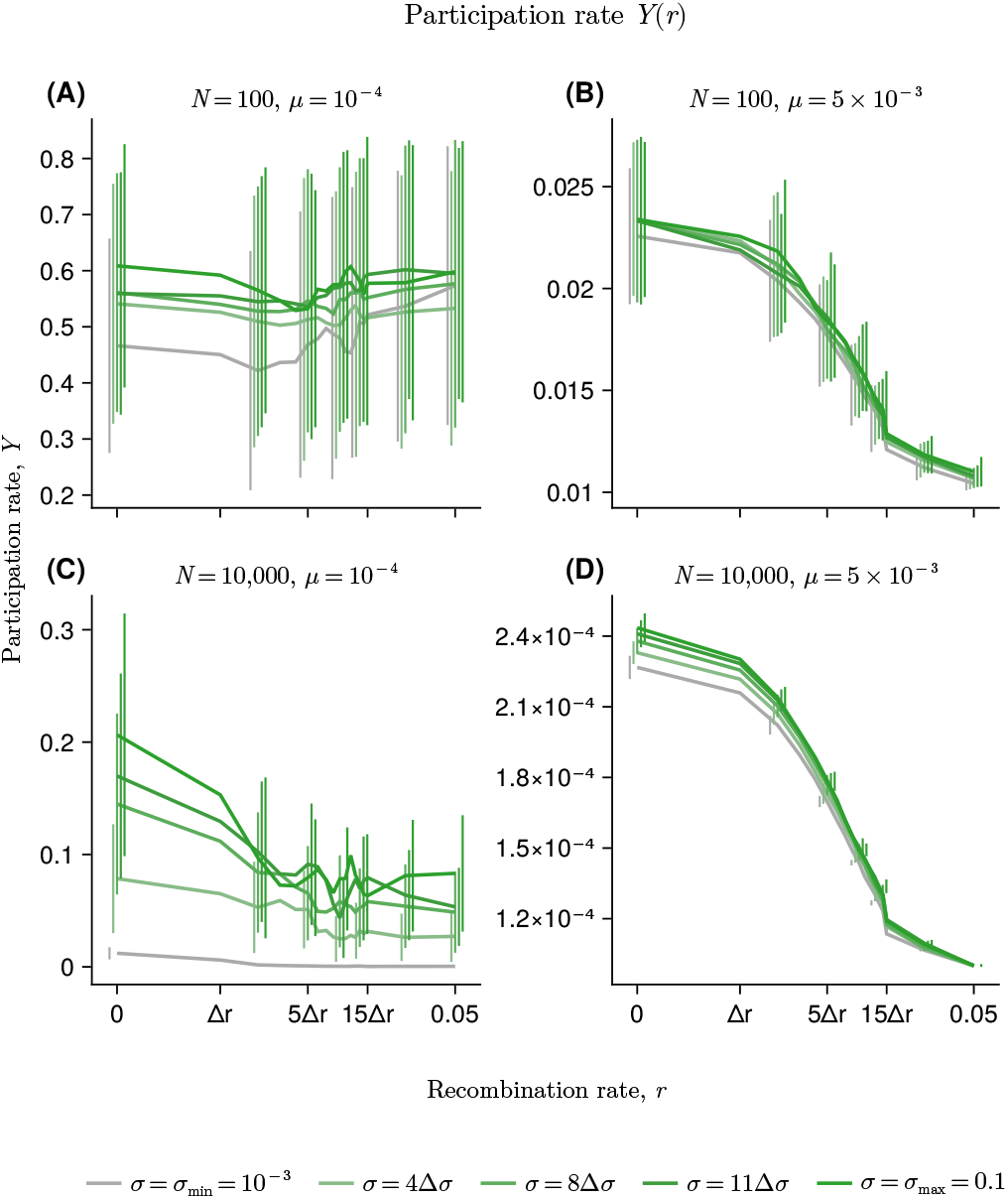
Effect of recombination rate on genotype concentration, quantified by the participation ratio 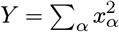, where *xα* is the frequency of genotype *α*. Each panel shows *Y* as a function of recombination rate *r* for a different combination of population size and mutation rate: *N* = 100, *µ* = 10^−4^ (**A**); *N* = 100, *µ* = 5 × 10^−3^ (**B**); *N* = 10^4^, *µ* = 10^−4^ (**C**); and *N* = 10^4^, *µ* = 5 × 10^−3^ (**D**). Lines and error bars show the mean and standard deviation across 30 replicate simulations. Darker green colors indicate larger values of *σ*, whereas gray colors indicate smaller values of *σ*. Overall, increasing recombination generally decreases *Y*, indicating reduced genotype condensation and an increased effective number of genotypes (see also the prior study that more quantitatively investigated the condensation of genotype using the metric *Y* (ref. ^39^), which used a different setup: starting from diverse genotypes under absence of mutations).

**Supplementary Fig. S4.**
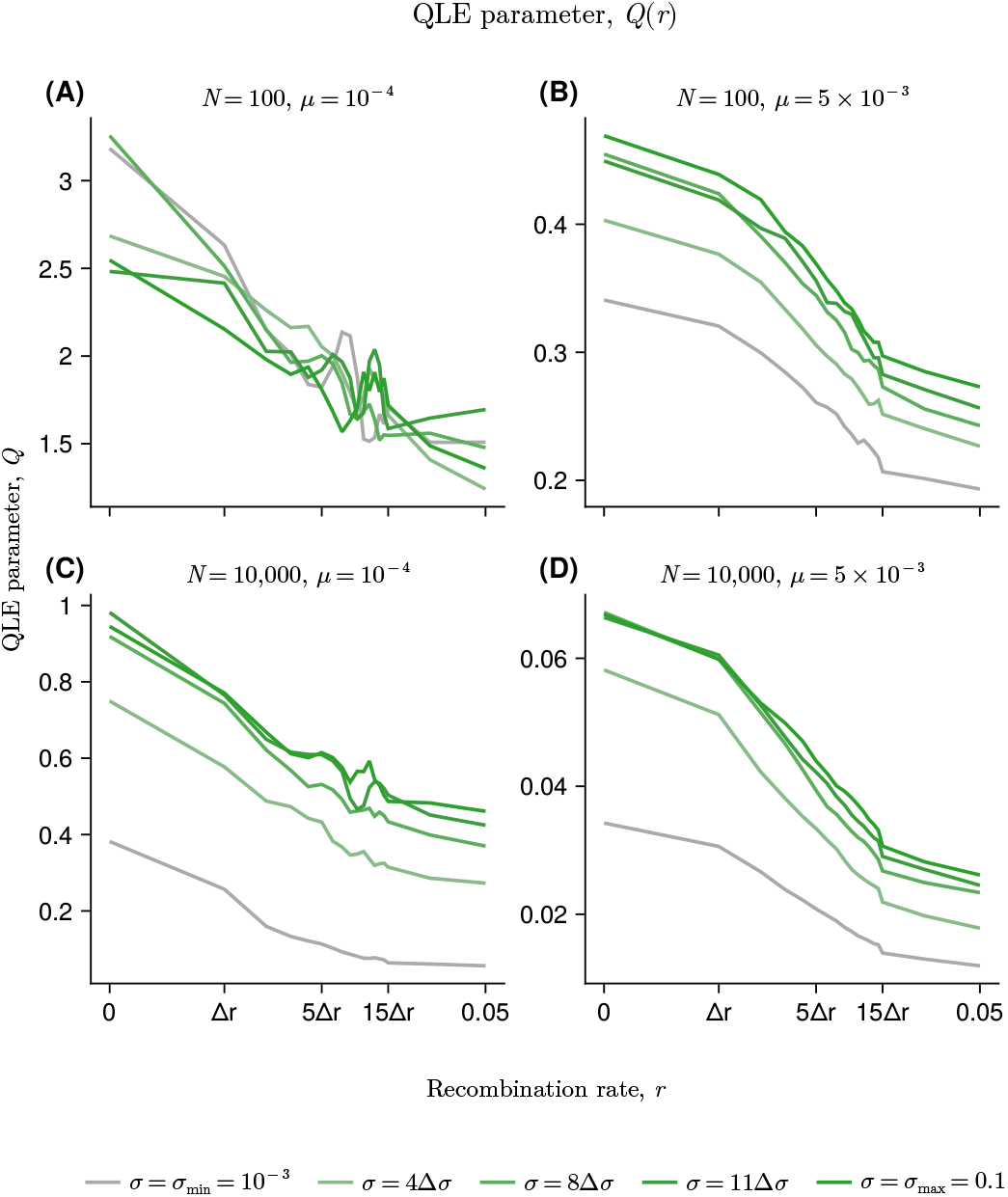
Effect of recombination rate on the QLE parameter *Q*, a metric that quantifies the average strength of pairwise associations among loci. Each panel shows *Q* as a function of recombination rate *r* for a different combination of *N* and *µ* (**A**-**D**). Lines and error bars show the mean and standard deviation across replicate simulations. Darker green colors indicate larger values of *σ*, whereas gray colors indicate smaller values of *σ*. As expected, frequent recombination generally reduces *Q* as recombination events break down the associations among loci. In contrast, increasing *σ* systematically increases *Q*, indicating that stronger selection or epistasis induces more nonrandom associations among loci.

**Supplementary Fig. S5.**
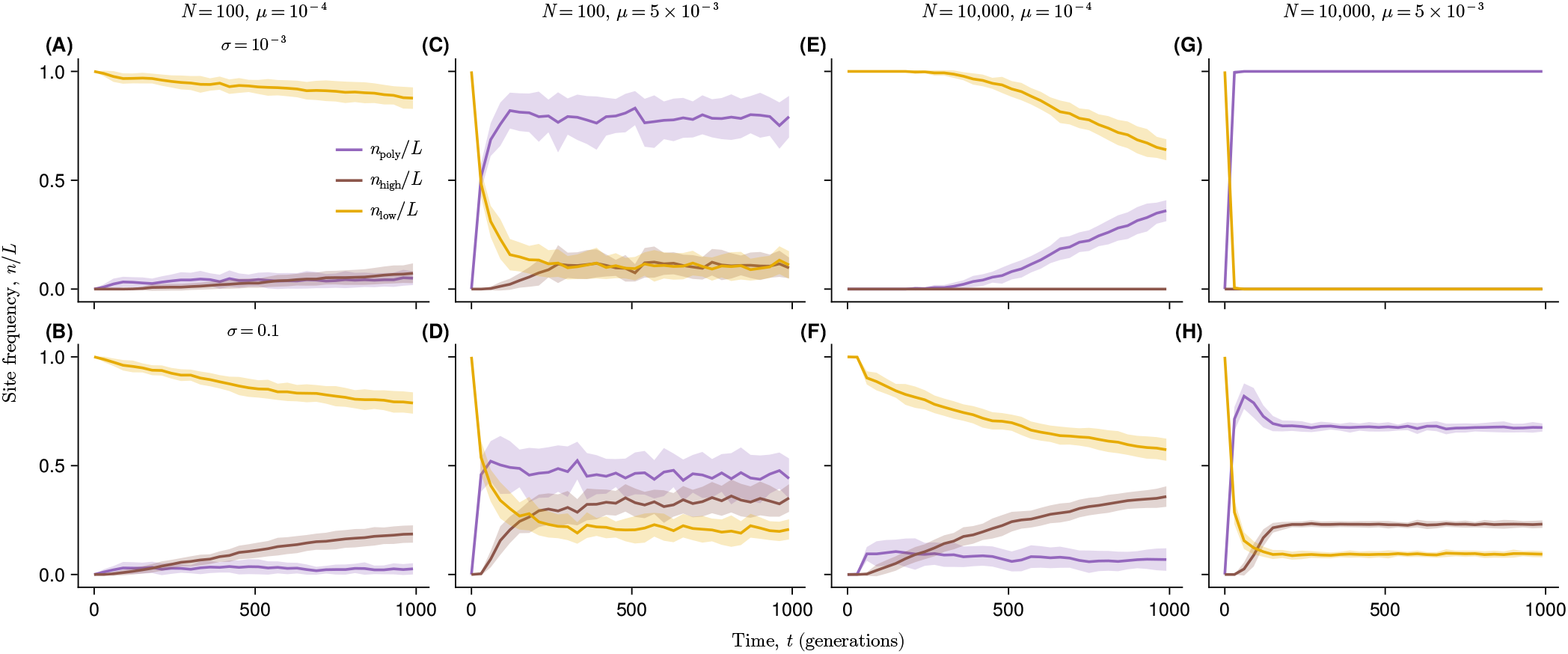
Here, *n*high and *n*low represent the number of sites at which the mutant allele frequency is above 90% or below 10%, respectively, and *n*poly denotes the number of polymorphic sites with mutant frequency between 10% and 90%. Thus, *n*high*/L, n*low*/L*, and *n*poly*/L* summarize the fractions of high-frequency, low-frequency, and polymorphic mutant sites, and sum to 1.0. Under weak selection (panels **A, C, E**, and **G**), mutant sites rarely reach high frequency, and larger mutation supply primarily increases *n*poly. When *µNL* is small, polymorphism remains limited, whereas larger *µNL* generates more polymorphic sites. Under strong selection (panels **B, D, F**, and **H**), *n*high increases and *n*low declines, reflecting the rise of beneficial mutations or high-fitness genetic backgrounds. With low mutation supply, the dynamics are sweep-like, whereas with high mutation supply, many mutations segregate simultaneously and the system approaches a dynamic balance between recurrent mutation and selection.

**Supplementary Fig. S6.**
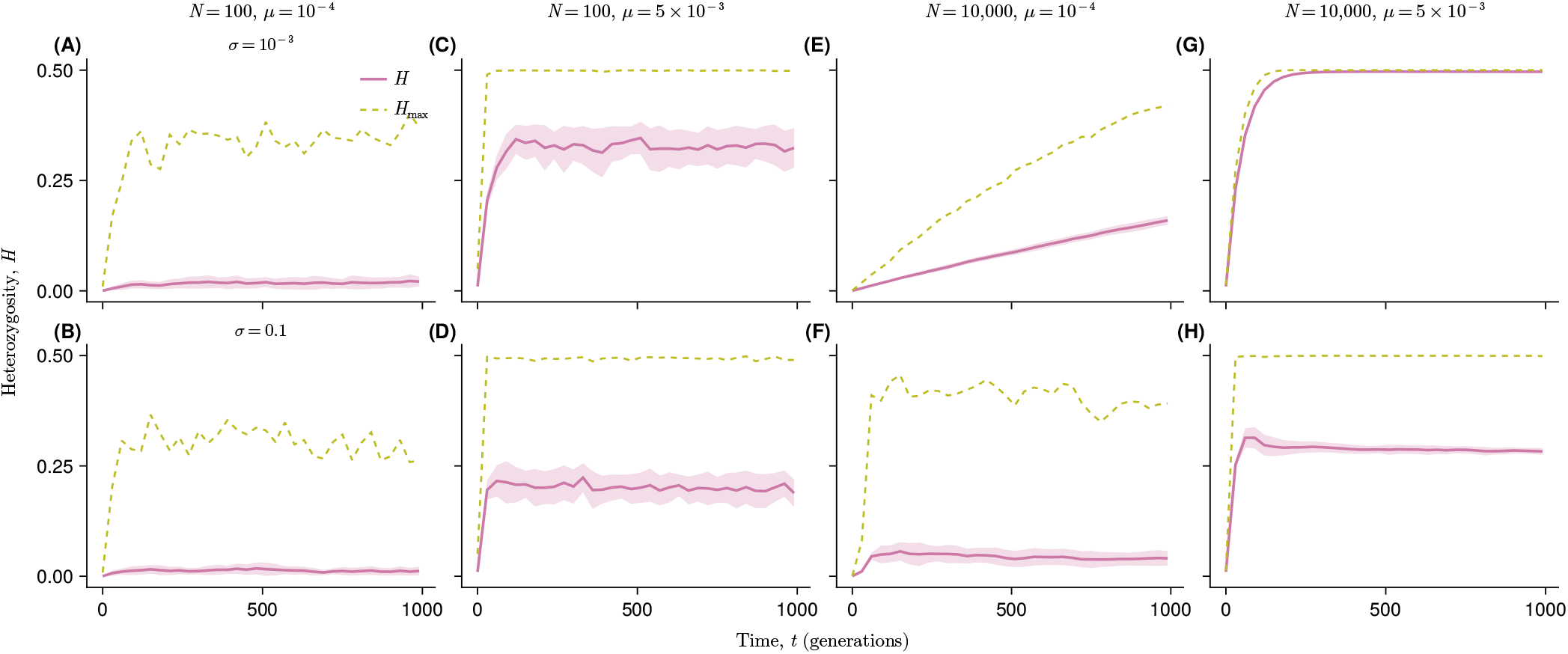
For small *σ*, evolution is close to neutral. The cases shown correspond to asexual evolution (*r* = 0). As expected, the site averaged heterozygosity *H* remains low when both population size and mutation rate are small (panel **A**). With larger *N* and low *µ, H* increases slowly and reaches moderate values (panel **E**). At high *µ, H* rises rapidly (panels **C** and **G**), and when *N* is also large, *H* approaches its maximum value. Under strong selection, the dynamics depend strongly on mutation supply. When both *N* and *µ* are small, mutation supply is limited and *H* remains near zero (panel **B**). When *µ* is high, even in the small population, *H* rises rapidly but fluctuates around intermediate values, reflecting recurrent mutation together with drift and stochastic lineage dynamics (panel **D**). With large *N* aand low *µ*, mutation supply is larger but still much lower than in the high-*µ* case (panel **F**). In this regime, average *H* remains low even though individual sites can transiently reach high heterozygosity as sweeping mutations pass through intermediate frequency before approaching fixation. Finally, with both large *N* and high *µ*, recurrent mutation maintains substantial polymorphism, while strong selection drives some sites toward high or low frequency (panel **H**). Thus, some sites remain near intermediate frequency, whereas others become nearly fixed, consistent with competition among selected genetic backgrounds.

**Supplementary Fig. S7.**
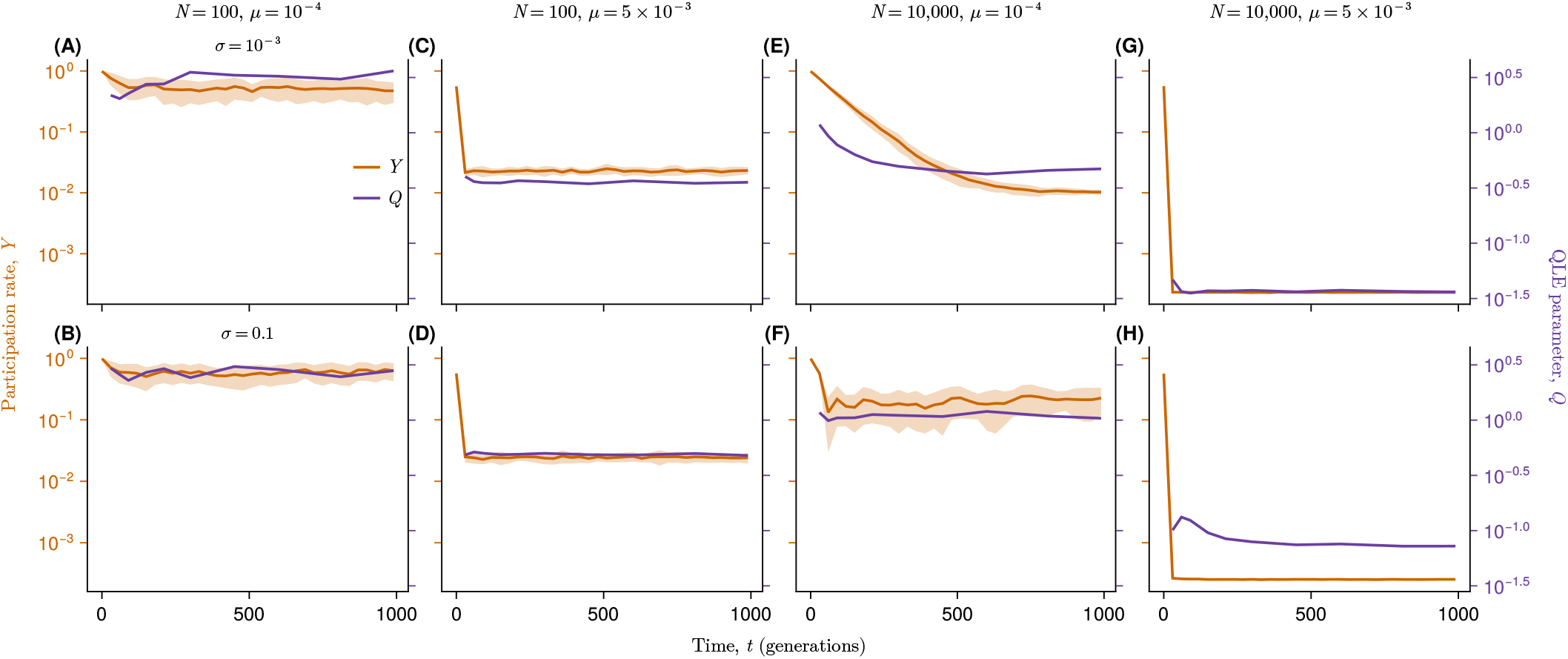
The cases shown correspond to *r* = 0 case. 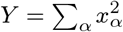, where *xα* is the frequency of genotype *α*, serves as a genotype-concentration parameter. Its inverse, 1*/Y*, represents the effective number of genotypes. We also define *Q* = Σ_*i<j*_ ⟨*χij* ⟩*/Σ* _*i<j*_ ⟨*χiiχjj* ⟩, where *χij* is the second-order cumulant and ⟨·⟩ denotes the average over independent simulations with the same underlying fitness parameters. Thus, *Q* measures LD relative to single-locus diversity. The difference is that *Y* measures concentration at the genotype level, whereas *Q* measures correlated allele structure across loci. These quantities are expected to behave similarly when genotype-level concentration creates linkage disequilibrium among many loci. Under neutral evolution, *Y* ∼ 1*/*(1 + 2*µNL*), and this approximation matches the steady-state values of *Y* for the weak-selection cases with *σ* = 10^−3^ (panels **A, C, E**, and **G**). Selection can increase *Y* by amplifying high-fitness genotypes; however, in our simulations, the effect of selection on *Y* was more subtle (panels **B, D, F**, and **H**). As expected, *Q* behaves similarly to *Y* . However, *Q* is generally higher in the larger-*σ* cases, even when *Y* is close to 1*/N* (for example, compare panels **G** and **H**). This suggests elevated nonrandom association between loci under stronger selection. Overall, *Y* and *Q* show qualitatively similar behavior, suggesting elevated LD is often associated with genotype concentration. However, elevated *Q* when *Y* is small suggests also that nonrandom associations can also persist even when the population is diversifed (*Y* ≃ 1*/N* ), likely due to selection and epistasis.

**Supplementary Fig. S8.**
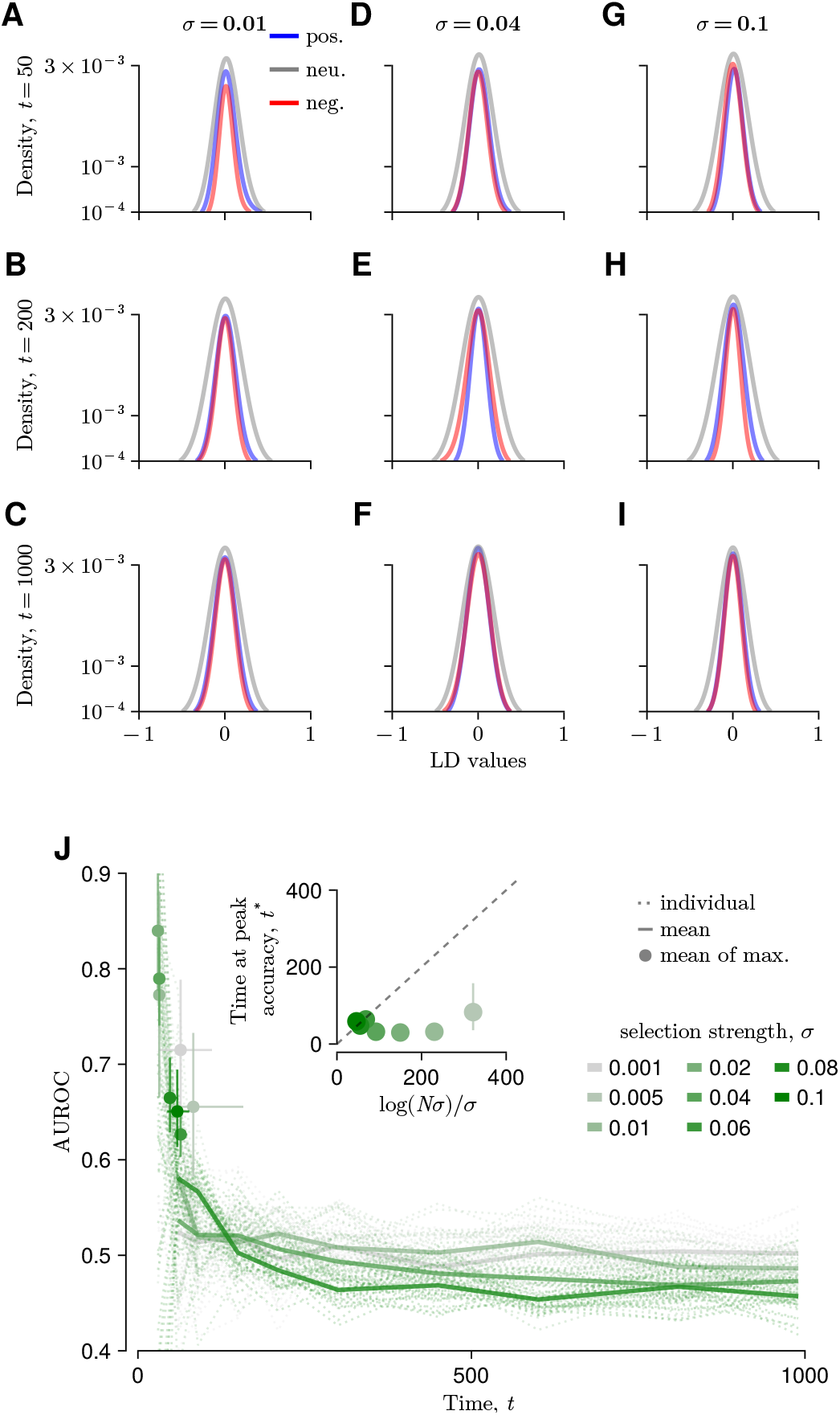
Distribution of LD for frequent recombination (*r* = 0.01) case. **A–I** Same analysis as **Fig. 3A-I**, but for the *N* = 10^3^ and *r* = 0.01 case rather than *r* = 0. Across all time points and *σ* values, the distributions collapse onto a normal distribution with zero mean. The AUROC value, used as accuracy metric, remains low across all times and *σ* values; therefore, unlike the *r* = 0 case, the peak time *t*^*^ is not simply related to log(*N σ*)*/σ* (**J**).

**Supplementary Fig. S9.**
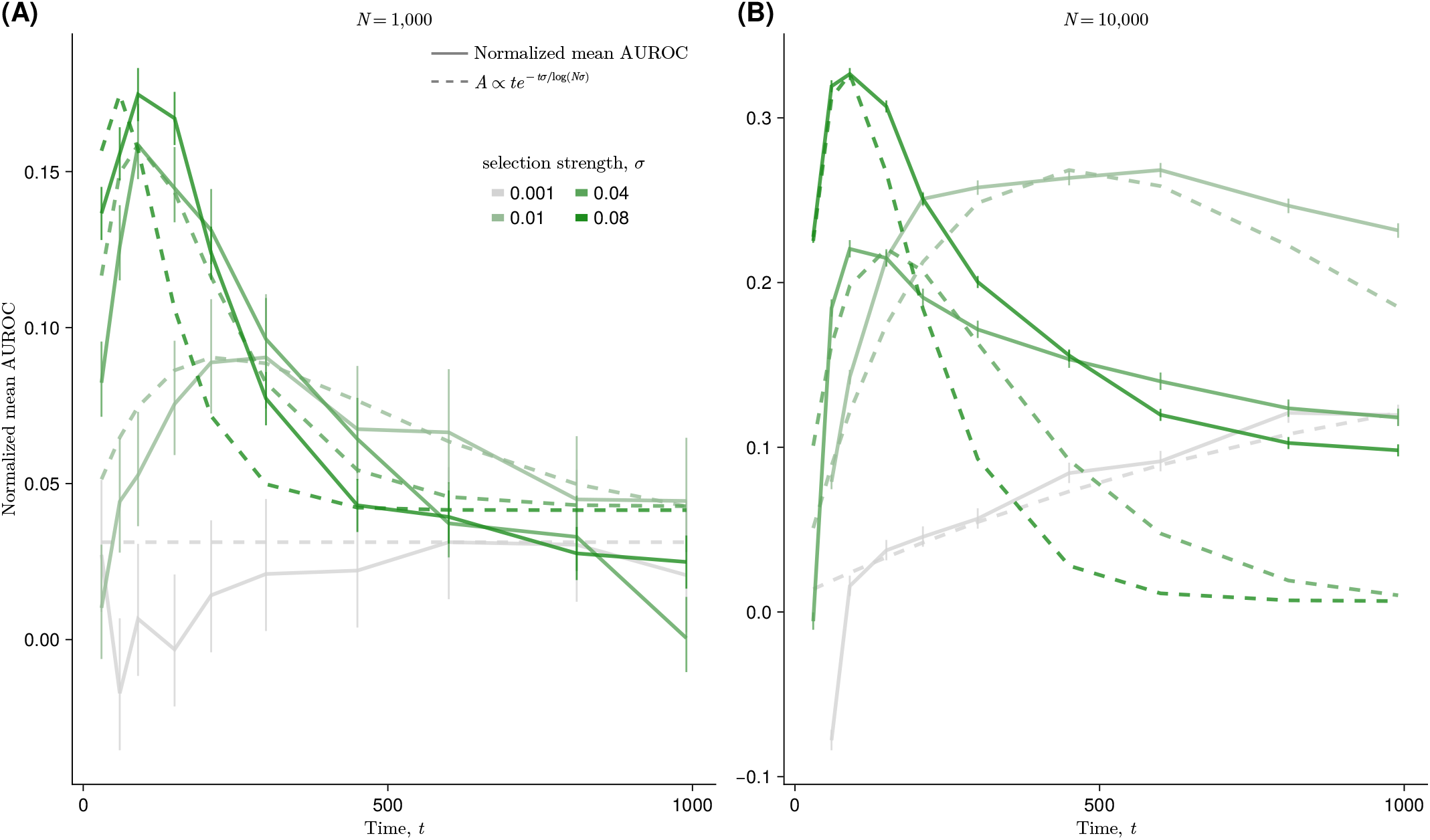
Fitting of the mean AUROC trajectories using the phenomenological expression of accuracy. Eq. (10) The presented mean AUROC trajectories are normalized by subtracting 0.5 from the raw AUROC values. The raw AUROC values are identical to those shown in **(A) Fig. 3J** and **(B) Fig. S11J** for the cases *N* = 10^3^ and *N* = 10^4^, respectively. The amplitude of the phenomenological accuracy function *A*(*t*) is estimated such that the peak values of *A*(*t*) and the observed mean values match. The overall profile of the accuracy trajectory, as well as the time at which the accuracy reaches its maximum, is well characterized by the phenomenological function. The error bars represent the 95% confidence intervals estimated across 100 inference results.

**Supplementary Fig. S10.**
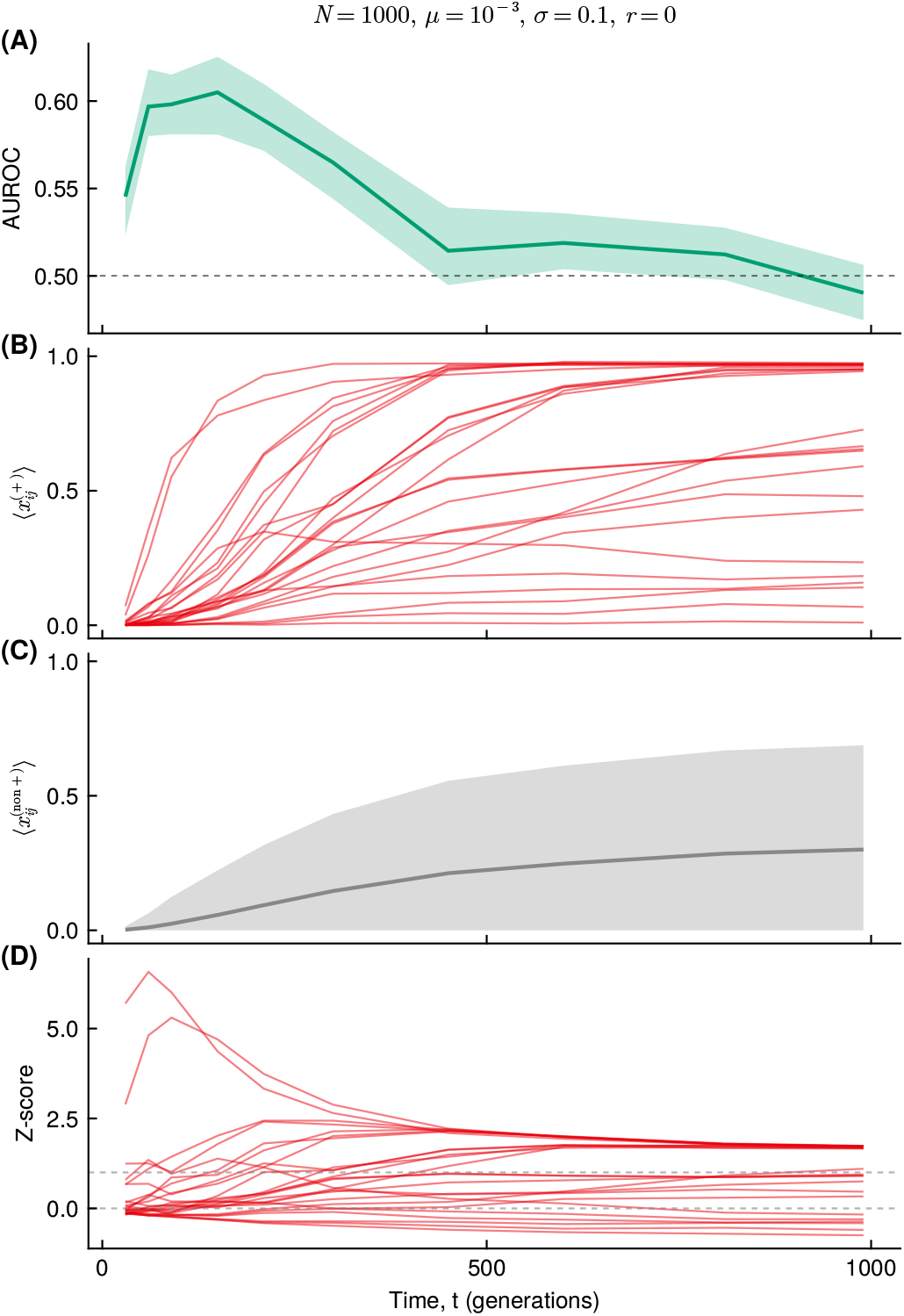
Dynamics of pairwise frequencies underlying the transient accuracy of LD-based epistasis detection. Results are shown for *N* = 10^3^, *µ* = 10^−3^, *r* = 0, and *σ* = 0.1. **(A)** AUROC for detecting positive epistatic interactions using the LD-based method, reproduced from **Fig. 4. (B)** Mean pairwise frequency of mutation pairs with positive epistasis, denoted 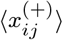. **(C)** Mean pairwise frequency of pairs without positive epistasis, denoted 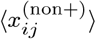. Solid lines and shaded regions indicate the mean and standard deviation across mutation pairs and 30 replicate simulations. **(D)** Standardized separation between positively epistatic pairs and the background, 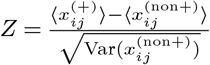, which quantifies how strongly positively epistatic pairs are enriched relative to non-positive pairs. The early increase in this separation explains the initial improvement in LD-based detection accuracy, whereas its later decrease reflects the accumulation of background linkage among non-positive pairs.

**Supplementary Fig. S11.**
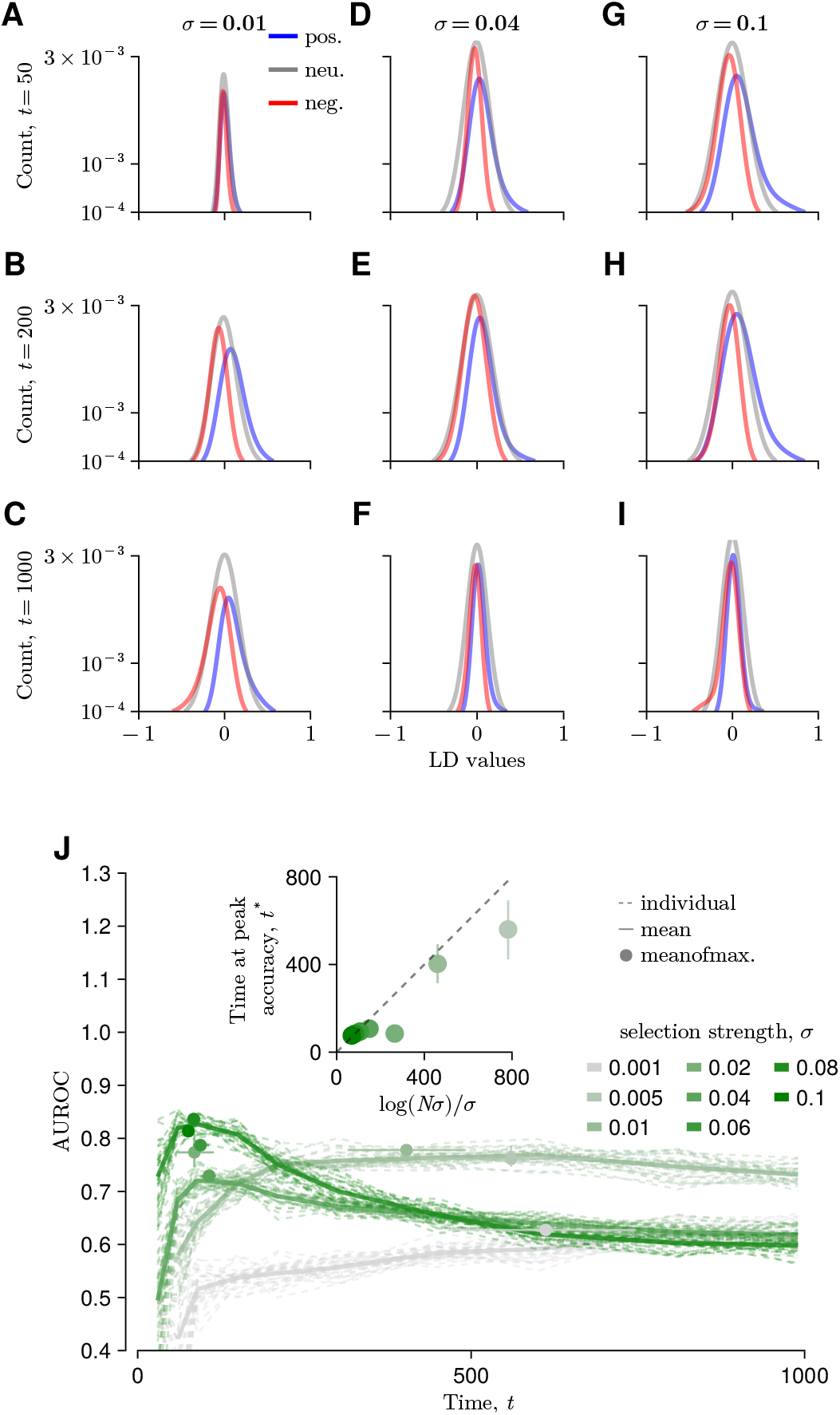
Distribution of LD for larger *N* case. **A–I** Distribution of LD values over time and selection strength *σ* for the *N* = 10^4^ and *r* = 0 case. The qualitative dependence on time and *σ* is the same as in the *N* = 10^3^ simulations: distributions become more separated at intermediate times and progressively overlap as time advances. The time window in which this separation appears also depends on *σ*. The average AUROC values based on LD show the same trend as in the *N* = 10^3^ case: as *σ* increases, the AUROC peak occurs earlier, and the peak time is approximately proportional to log(*N σ*)*/σ* (**J**).

**Supplementary Fig. S12.**
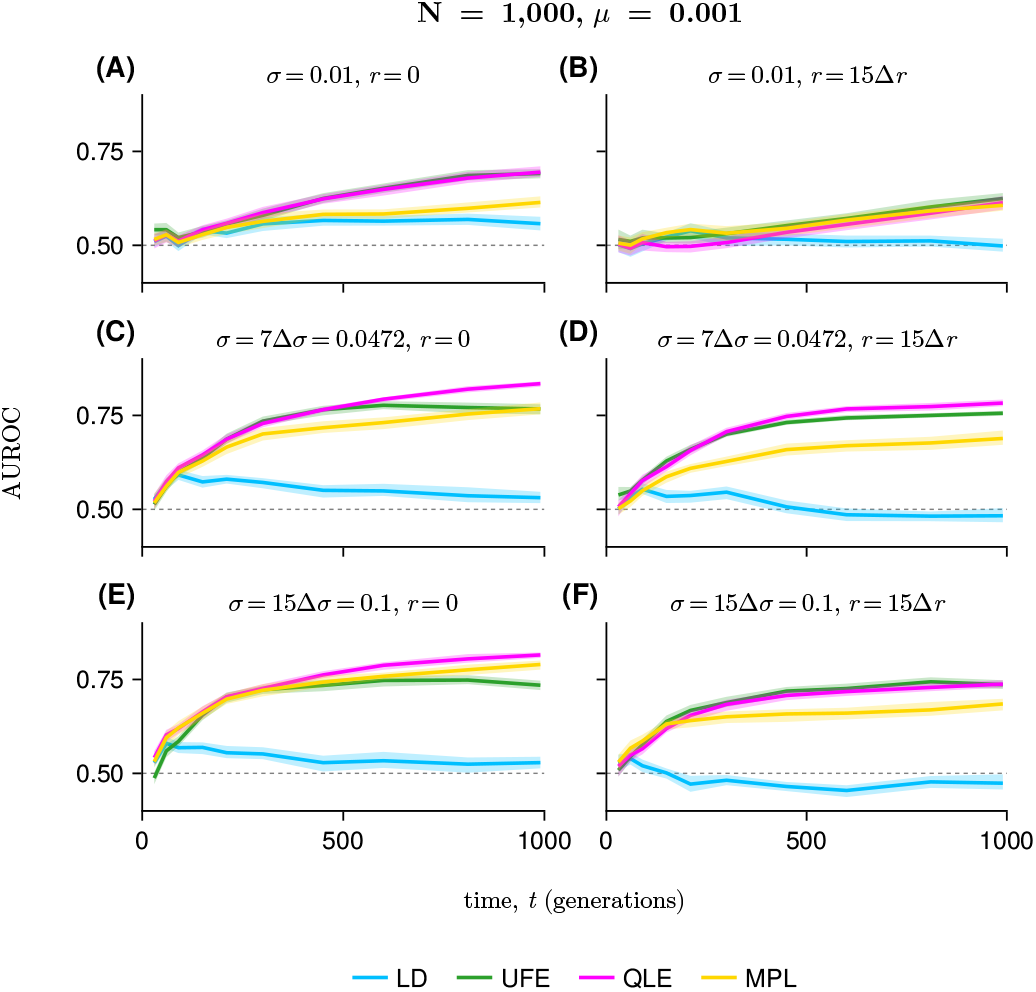
Detection accuracy for negative epistatic interactions as a function of time. This figure corresponds to **Fig. 4**, which is for the positive epistasis case. Population size and mutation rate are fixed at *N* = 1000 and *µ* = 10^−3^. The left column shows simulations without recombination, *r* = 0, and the right column shows simulations with high recombination frequency, *r* = 0.01. Overall, the qualitative patterns are similar to those observed for positive epistasis: accuracy is lower under weak selection, increases with selection strength, and is generally reduced by recombination. AUROC values are slightly lower for negative epistasis than for positive epistasis cases.

**Supplementary Fig. S13.**
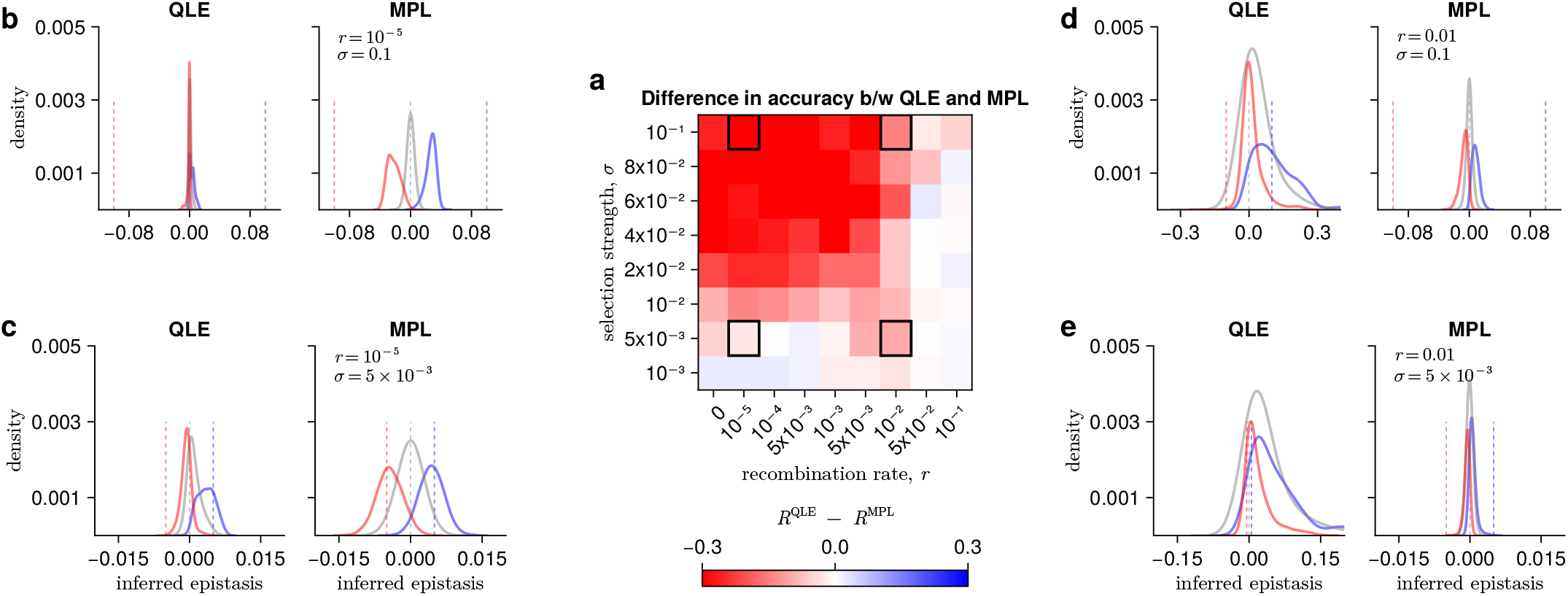
Results of inferred epistasis from WF simulations with *N* = 10^4^ and *µ* = 10^−3^ using the MPL and QLE methods, visualized through the difference in Pearson’s *R* between MPL and QLE (*R*^QLE^ − *R*^MPL^). In this larger-*N* setting, MPL outperforms QLE across most parameter regimes. The distributions for positive, negative, and neutral epistasis are more clearly separated, especially in the low-recombination regime, when using MPL.

**Supplementary Fig. S14.**
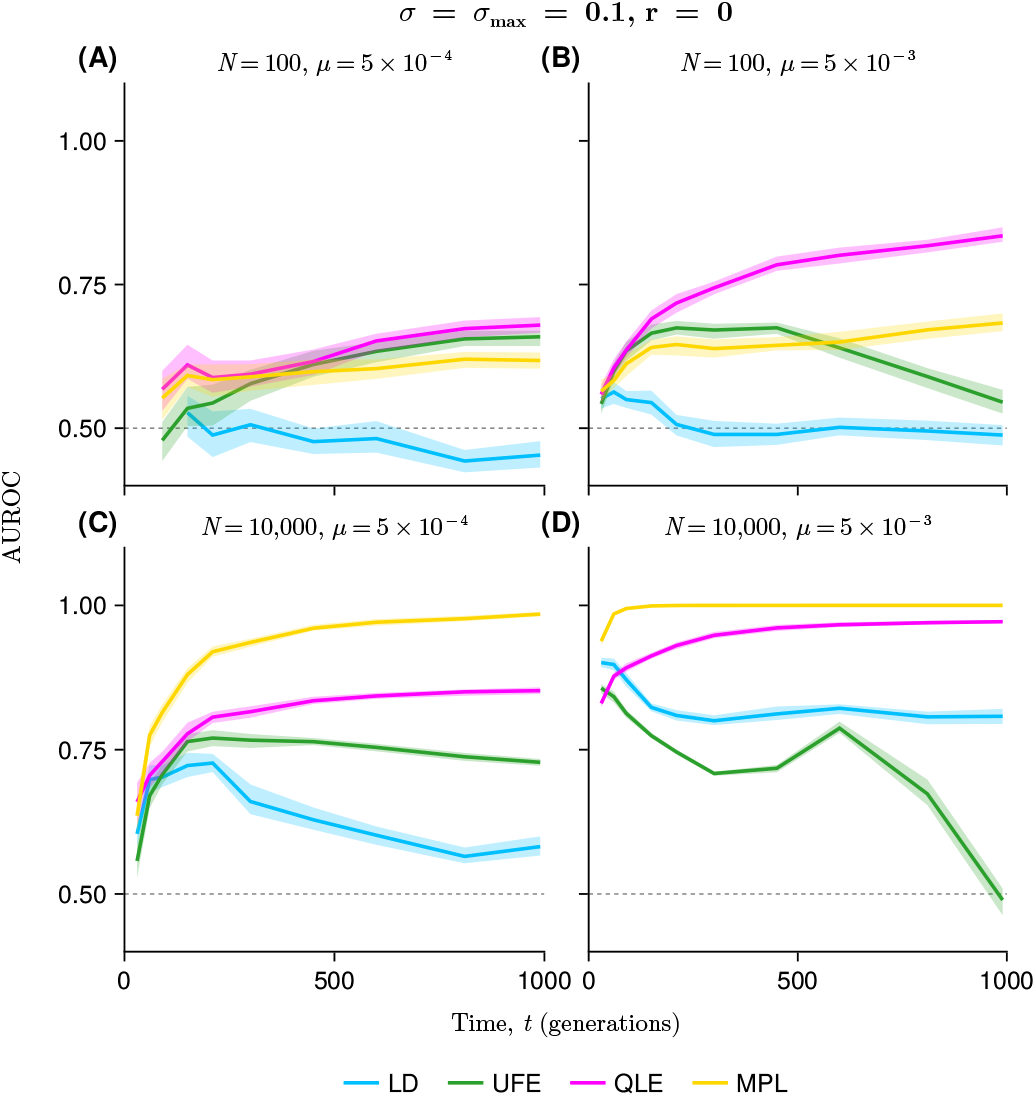
Time dependence of epistasis inference accuracy for representative combinations of population size and mutation rate under strong selection without recombination. Here, *σ* = 0.1 and *r* = 0. Each panel shows AUROC over time for the LD, UFE, QLE, and MPL methods. Panels correspond to *N* = 100, *µ* = 5 × 10^−4^ (**A**); *N* = 100, *µ* = 5 × 10^−3^ (**B**); *N* = 10^4^, *µ* = 5 × 10^−4^ (**C**); and *N* = 10^4^, *µ* = 5 × 10^−3^ (**D**). The relative performance of QLE and MPL depends on population size *N* . In smaller populations, QLE tends to reach higher accuracy than MPL, whereas in larger *N*, MPL generally outperforms QLE. At high *µ* and large *N* values, QLE and MPL become more similar in performance. UFE shows a distinct behavior; its accuracy can decrease over time, especially at high *µ*.

**Supplementary Fig. S15.**
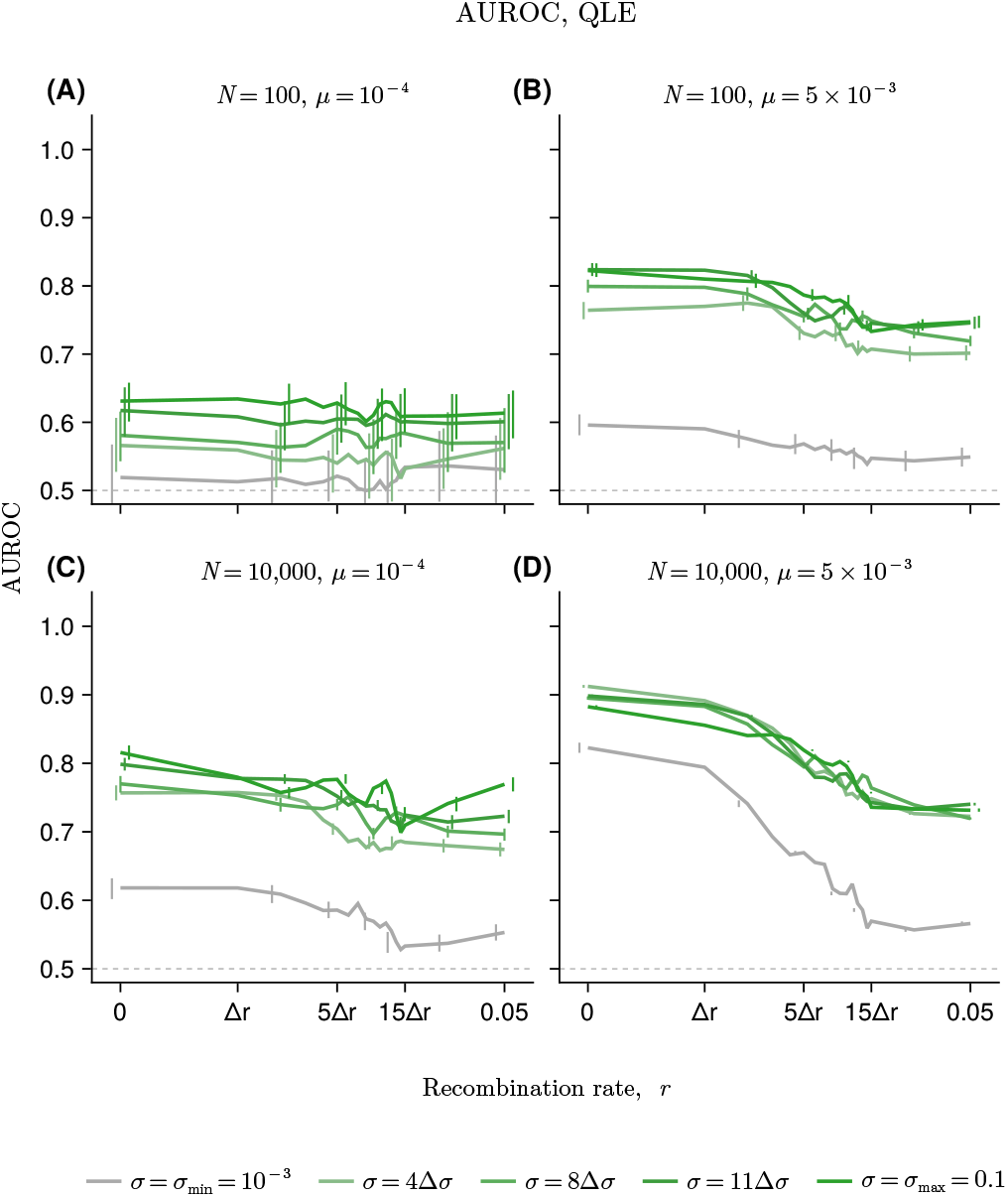
Effect of recombination rate on epistasis-inference accuracy for the QLE method. Each panel shows AUROC as a function of recombination rate *r* for *N* = 100, *µ* = 10^−4^ (**A**); *N* = 100, *µ* = 5 × 10^−3^ (**B**); *N* = 10^4^, *µ* = 10^−4^ (**C**); and *N* = 10^4^, *µ* = 5 × 10^−3^ (**D**). Lines and error bars show the mean and standard deviation across 30 replicate simulations. Across parameter regimes, QLE accuracy generally decreases as recombination rate increases, consistent with the reduction of associations among loci. An increase in the recombination decreases the accuracy and eventually the AUROC values from *N* = 100 and *N* = 10^4^ become comparable at the highest *r* value, even though the AUROC value is significantly higher for the *N* = 10^4^ case than the *N* = 100 case at *r* = 0. We also confirmed that the recombination does not impact the accuracy of selection inference.

**Supplementary Fig. S16.**
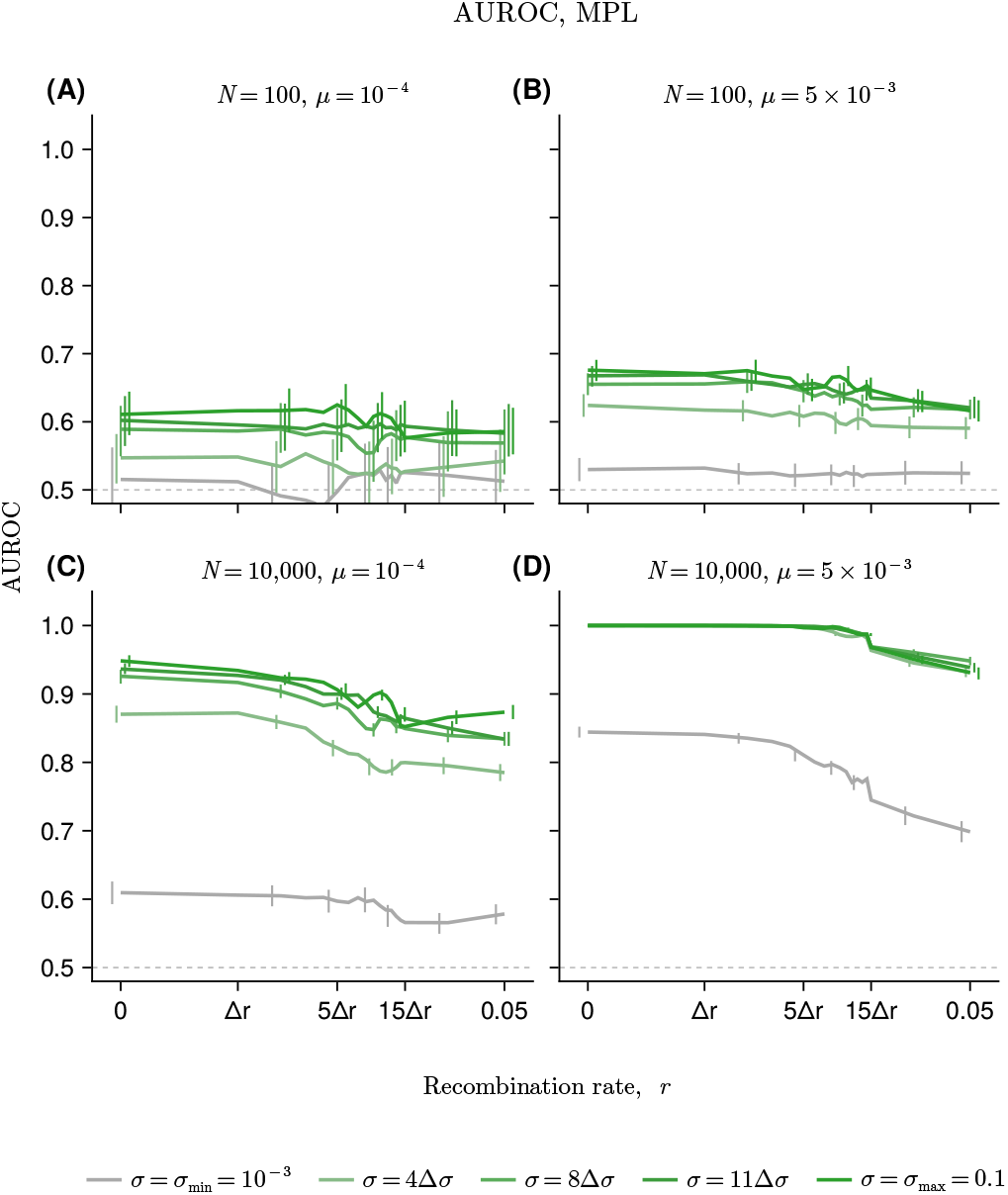
Effect of recombination rate on epistasis-inference accuracy for the MPL method. MPL accuracy typically depends on the *σ* value; a higher *σ* value results in a higher AUROC value. Recombination can also reduce accuracy for the MPL case.

**Supplementary Fig. S17.**
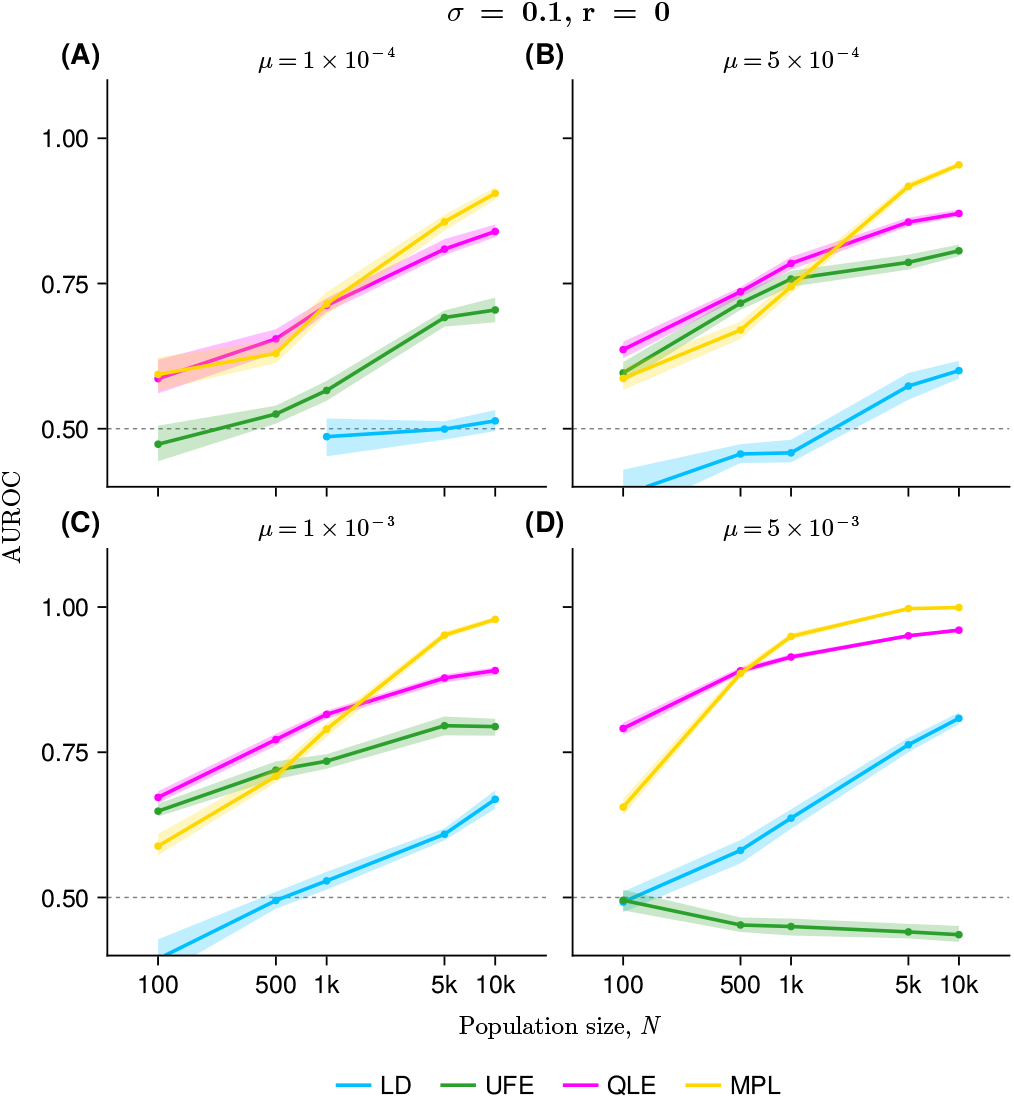
Effect of population size and mutation rate on detection accuracy for negative epistatic interactions. This figure corresponds to **Fig. 7**, which is the case for positive epistasis. Here, *σ* = 0.1 and *r* = 0. The population-size dependence is also qualitatively similar to the positive-epistasis case. Accuracy generally increases with *N* . The relative performance of the methods is also largely conserved, with QLE performing relatively well in smaller populations and MPL improving more strongly in larger populations. Absolute AUROC values are slightly lower than for positive epistasis overall.

**Supplementary Fig. S18.**
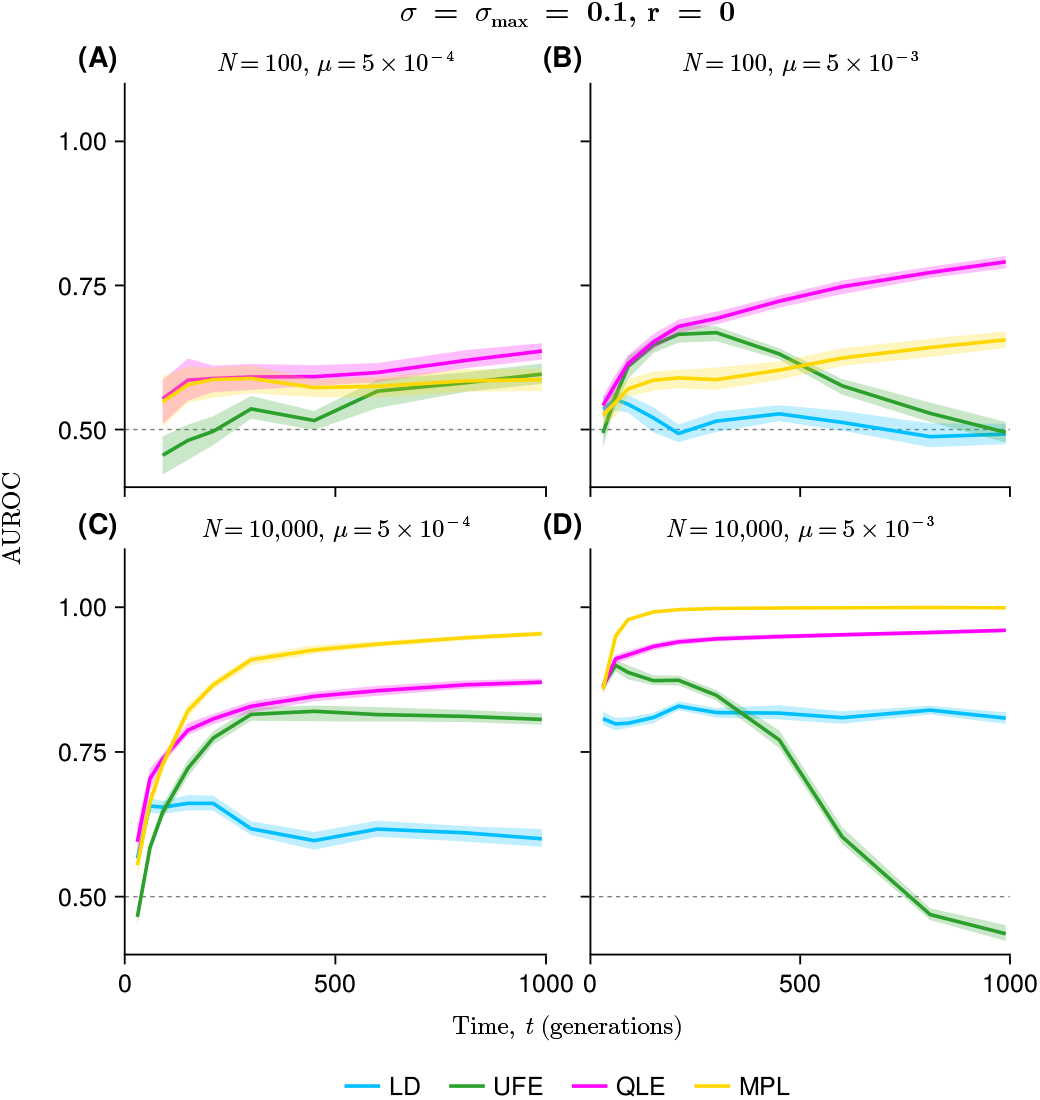
Time dependence of the detection accuracy for negative epistasis under representative *N* and *µ* combinations. This result corresponds to **Fig. S14**, which is for positive epistasis. The LD result is not shown for the *N* = 100, *µ* = 5 ×10^−4^ case because no detectable mutation pairs passed the pair-frequency threshold used for the LD-based analysis. Overall, the time-dependent patterns and relative performance of the methods are similar to those observed for positive epistasis, although the absolute AUROC values are slightly lower. A notable difference is that the early-time peak characteristic of LD-based detection for positive epistasis becomes unclear for negative epistasis.

**Supplementary Fig. S19.**
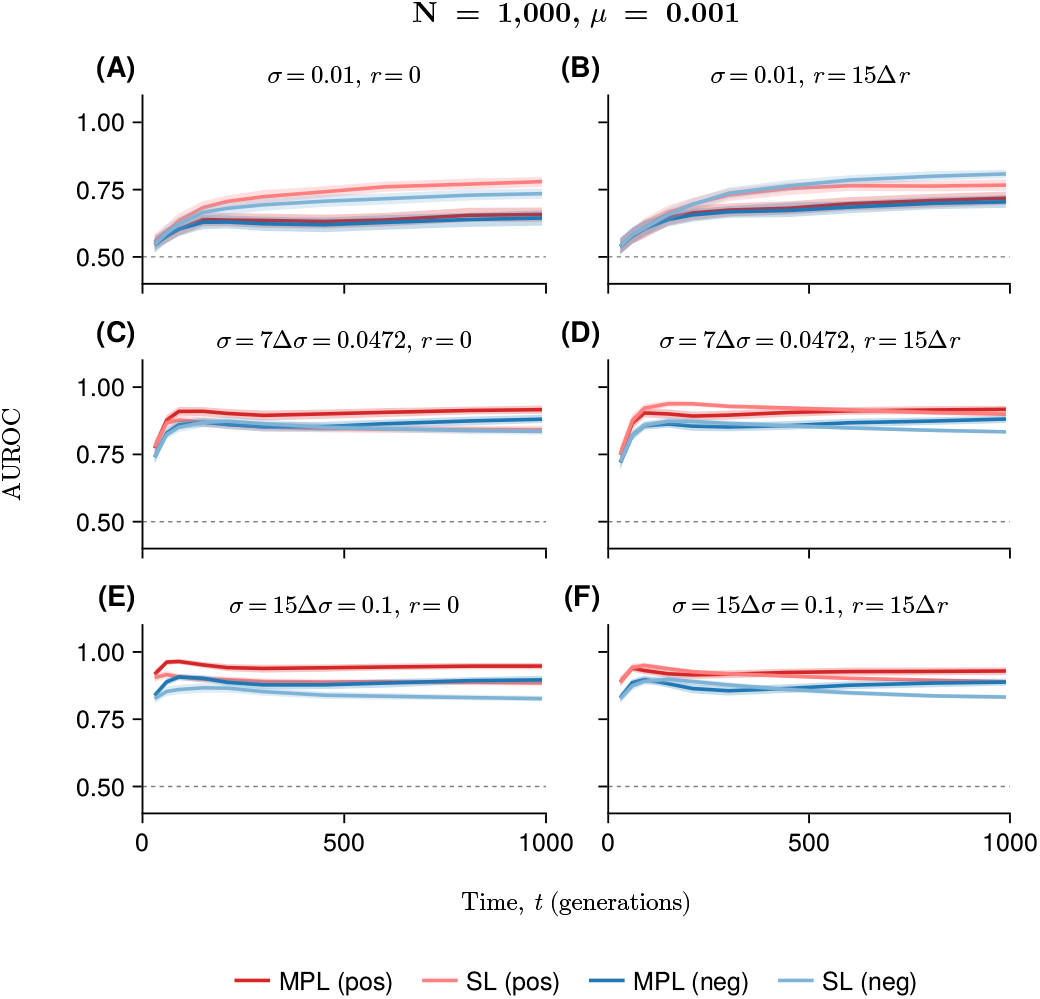
Selection-inference accuracy over time. Population size and mutation rate are fixed at *N* = 1000 and *µ* = 10^−3^. Here we focused on asexual (left) and frequent recombination (right) cases. The mean (bold line) and standard deviation (shaded ribbon) values are estimated from 30 replicates. Selection inference becomes more accurate as selection strength increases, consistent with the epistasis inference case. Typically AUROC values rapidly increase and remain high over a broad time window. Recombination has little effect on the accuracy. MPL generally outperforms the single-locus (SL) estimator ^40^, and positive selection is inferred more accurately than negative selection.

**Supplementary Fig. S20.**
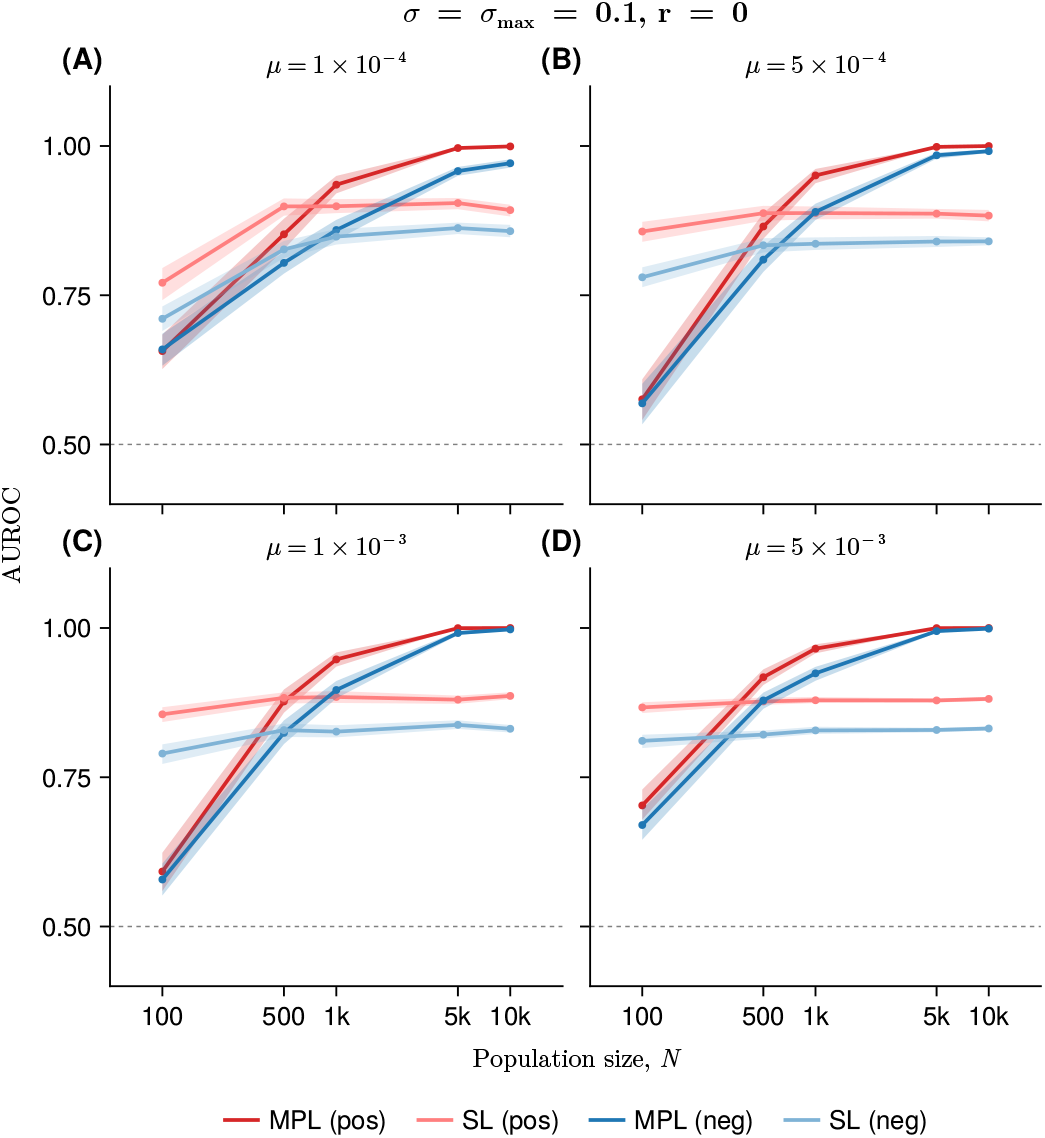
Effect of population size and mutation rate on selection-inference accuracy. AUROC values are computed at *t* = 1000. The SL estimator shows relatively weak dependence on population size *N* and tends to plateau at intermediate AUROC values. In contrast, MPL improves with increasing *N*, reaching 1.0 vlaue in large populations. The same qualitative pattern is observed across mutation rates).

**Supplementary Fig. S21.**
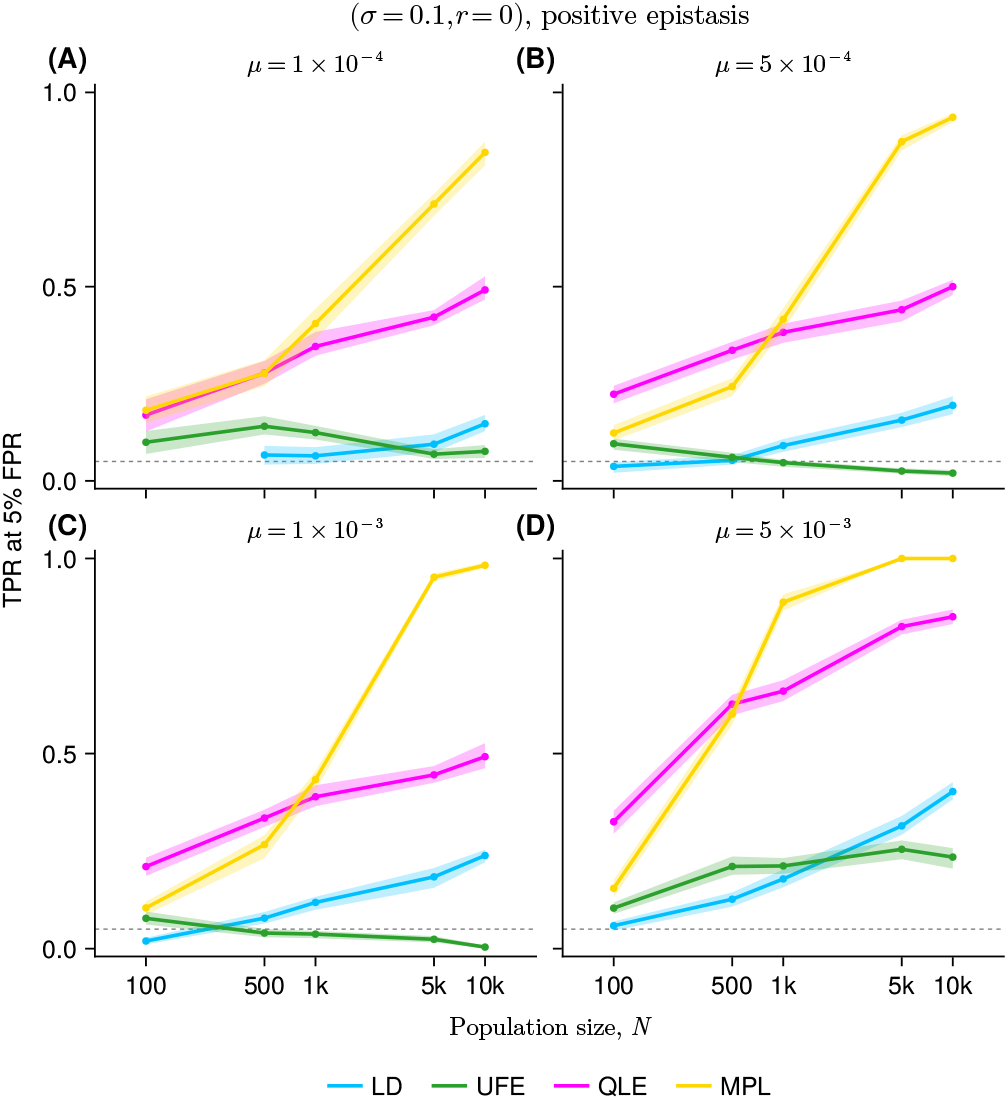
True-positive rate at a fixed false-positive rate as a measure of statistical power for epistasis detection. The true-positive rate at 5% false-positive rate (TPR@5% FPR) is shown as a function of population size for four mutation-rate conditions. Selection strength and recombination rate are fixed at *σ* = 0.1 and *r* = 0, respectively. Each curve shows the mean TPR@5% FPR across 30 replicate simulations, and shaded ribbons indicate the standard deviation. Consistent with the AUROC-based analysis in **Fig. 7**, inference accuracy generally increases with *N* and *µ*, particularly for QLE- and MPL-based approaches.

**Supplementary Fig. S22.**
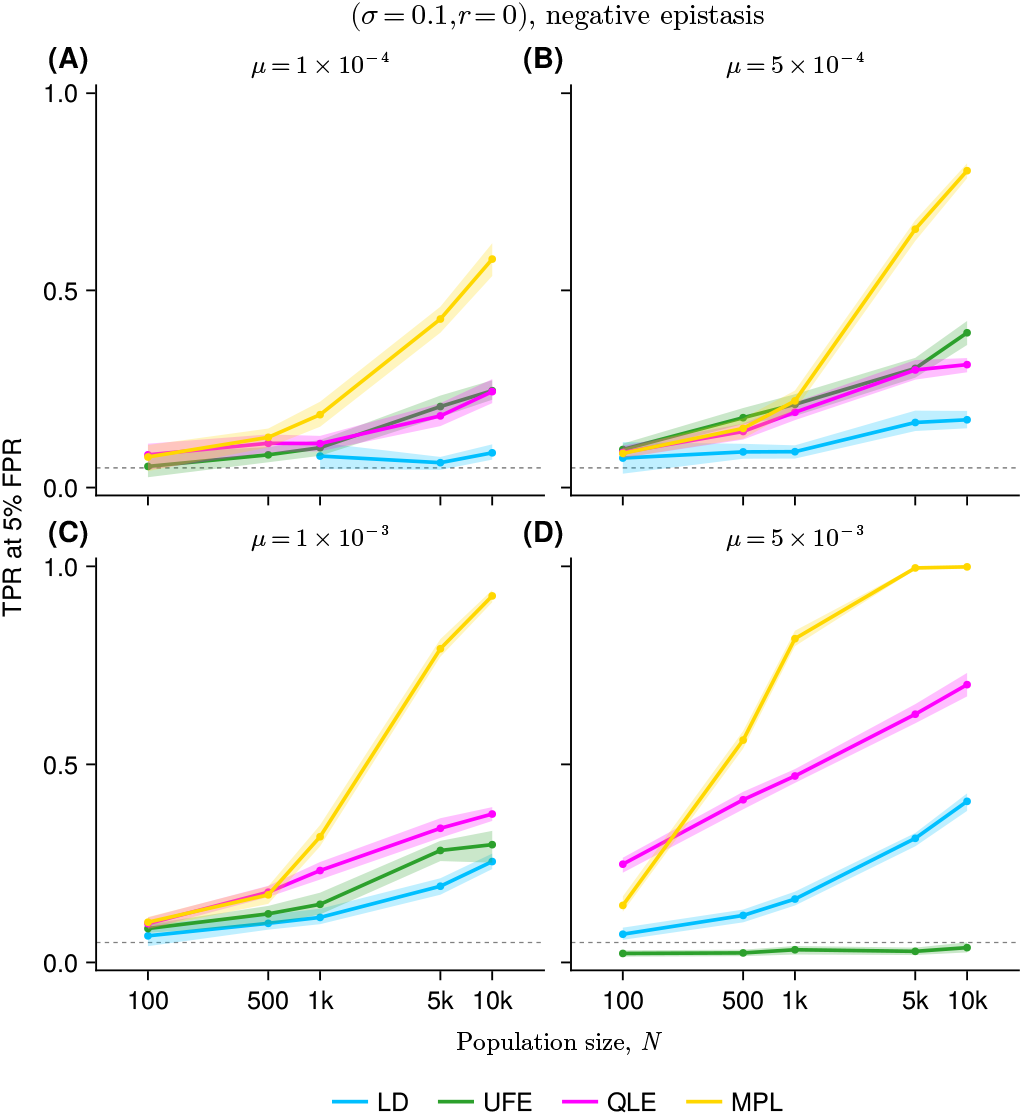
TPR@5% FPR is shown for detecting negative epistatic interactions as a function of population size *N* across multiple mutation rates *µ*. Although the absolute TPR@5% FPR values tend to be lower than those for positive epistasis, they are largely consistent with the positive epistasis case (**Supplementary Figure S21**).

**Supplementary Fig. S23.**
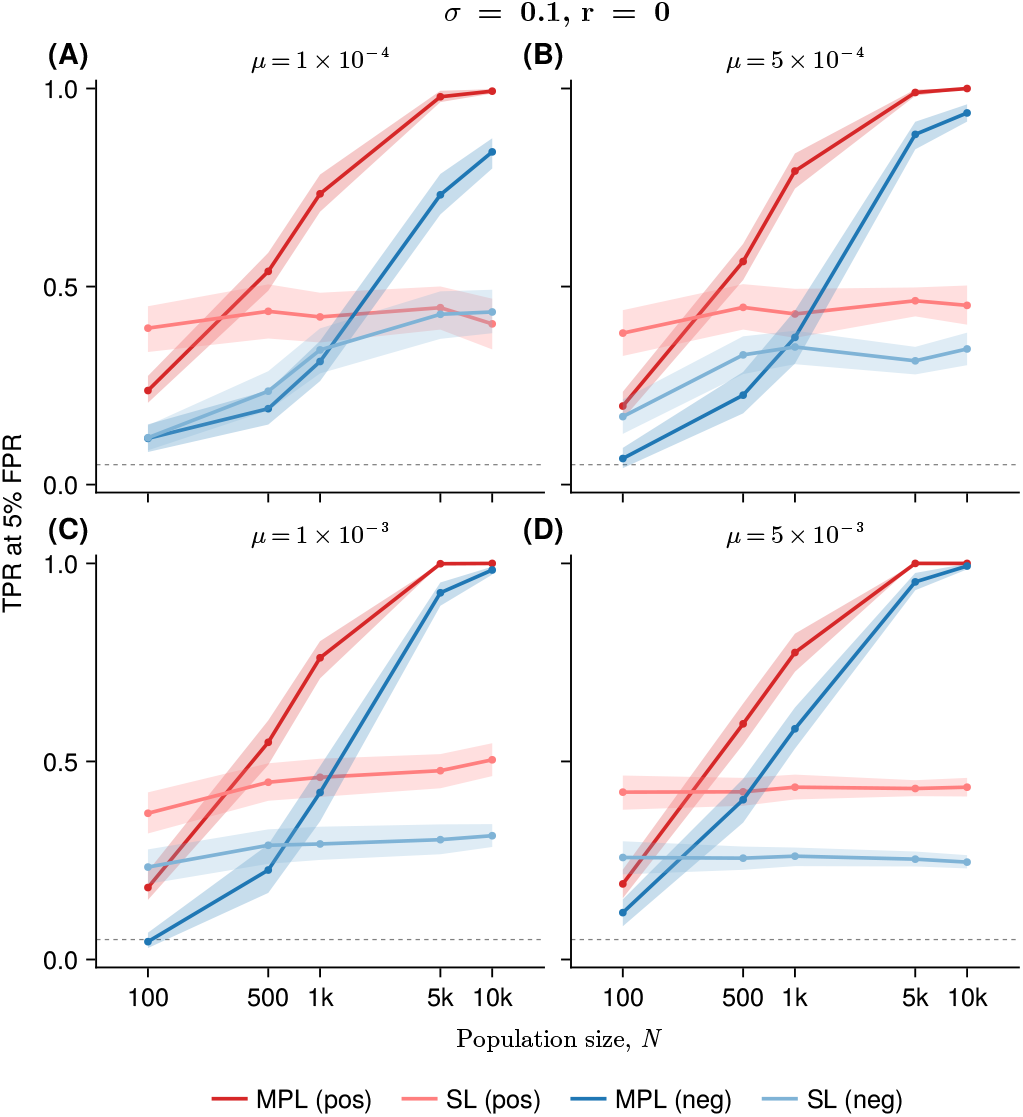
TPR@5% FPR for detecting positive and negative selection coefficients. The selection strength and recombination rate are fixed at *σ* = 0.1 and *r* = 0. Overall patterns are consistent with the AUROC-based analysis, but the difference between positive and negative cases is more pronounced under the TPR@5% FPR metric than under AUROC (**Supplementary Figure S20**).

**Supplementary Table 1.** Kolmogorov-Smirnov (KS) test *D* statistics for LD distributions, *N* = 10^3^, *r* = 0 (Fig. **3**). The times that achieve the maximum *D* statistics among three time points are shown in bold. The subcolumns labeled pos/neg, neg/neu, and pos/neg represent the comparisons of the density of LD values for positive versus neutral epistasis, negative versus neutral, and positive versus negative, respectively. In the case of smaller selective pressure (*σ* = 0.01), the peak time occurs later. In contrast, for higher values of *σ* = 0.1, the peak time often appears earlier. For an intermediate selection strength of *σ* = 0.04, the peak time is slightly later. Overall, increases in *σ* lead to earlier maximum *D* values and greater separation of the distributions.

| $t$ | $\sigma = 0.01$ | | | $\sigma = 0.04$ | | | $\sigma = 0.10$ | | |
| --- | --- | --- | --- | --- | --- | --- | --- | --- | --- |
|  | pos/neu | neg/neu | pos/neg | pos/neu | neg/neu | pos/neg | pos/neu | neg/neu | pos/neg |
| 50 | 0.051 | 0.053 | 0.082 | 0.120 | 0.118 | 0.230 | **0.221** | 0.137 | **0.353** |
| 200 | **0.125** | **0.106** | **0.225** | **0.200** | **0.144** | **0.328** | 0.145 | **0.167** | 0.297 |
| 1000 | 0.116 | 0.102 | 0.183 | 0.157 | 0.106 | 0.239 | 0.145 | 0.125 | 0.251 |

**Supplementary Table 2.** KS test *D* statistics for LD distributions, *N* = 10^4^, *r* = 0. Overall, the trends observed are similar to those seen in the small population size scenario. However, the time at which the highest *D* values are achieved occurs earlier among the three time points. The *D* values for the cases with a population size of *N* = 10, 000 are generally larger than those for the smaller population sizes (**Supplementary Table 1**), suggesting an enhanced separation of distributions in larger populations.

| $t$ | $\sigma = 0.01$ | | | $\sigma = 0.04$ | | | $\sigma = 0.10$ | | |
| --- | --- | --- | --- | --- | --- | --- | --- | --- | --- |
|  | pos/neu | neg/neu | pos/neg | pos/neu | neg/neu | pos/neg | pos/neu | neg/neu | pos/neg |
| 50 | 0.091 | 0.090 | 0.175 | **0.315** | 0.222 | **0.502** | **0.406** | **0.318** | **0.623** |
| 200 | **0.376** | 0.310 | **0.635** | 0.270 | **0.232** | 0.480 | 0.353 | 0.284 | 0.559 |
| 1000 | 0.327 | **0.336** | 0.567 | 0.207 | 0.193 | 0.397 | 0.152 | 0.229 | 0.373 |

**Supplementary Table 3.**
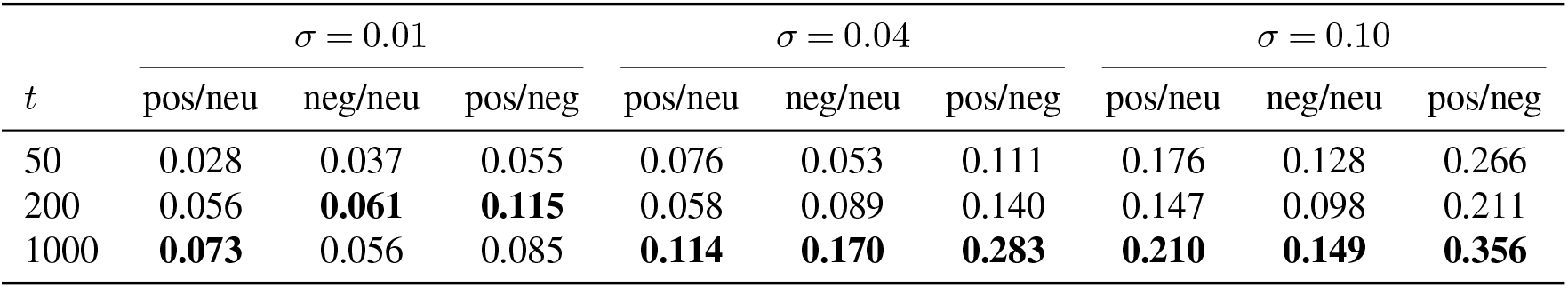
KS test *D* statistics for LD distributions, *N* = 10^3^, *r* = 0.01. In contrast to cases where recombination is absent (**Supplementary Tables 1-2**), the observed pattern in asexual evolution (increasing sigma leads to an earlier separation of the LD distributions of positive/neutral, negative/neutral, and positive/negative) becomes ambiguous; the time at which the maximum *D* values are reached occurs later in evolution overall. Additionally, *D* values tend to be lower than the asexual case (**Supplementary Table 1**) in the presence of recombination, particularly under weak selection.

| $t$ | $\sigma = 0.01$ | | | $\sigma = 0.04$ | | | $\sigma = 0.10$ | | |
| --- | --- | --- | --- | --- | --- | --- | --- | --- | --- |
|  | pos/neu | neg/neu | pos/neg | pos/neu | neg/neu | pos/neg | pos/neu | neg/neu | pos/neg |
| 50 | 0.028 | 0.037 | 0.055 | 0.076 | 0.053 | 0.111 | 0.176 | 0.128 | 0.266 |
| 200 | 0.056 | **0.061** | **0.115** | 0.058 | 0.089 | 0.140 | 0.147 | 0.098 | 0.211 |
| 1000 | **0.073** | 0.056 | 0.085 | **0.114** | **0.170** | **0.283** | **0.210** | **0.149** | **0.356** |

